# TINNiK: Inference of the Tree of Blobs of a Species Network Under the Coalescent

**DOI:** 10.1101/2024.04.20.590418

**Authors:** Elizabeth S. Allman, Hector Baños, Jonathan D. Mitchell, John A. Rhodes

## Abstract

The tree of blobs of a species network shows only the tree-like aspects of relationships of taxa on a network, omitting information on network substructures where hybridization or other types of lateral transfer of genetic information occur. By isolating such regions of a network, inference of the tree of blobs can serve as a starting point for a more detailed investigation, or indicate the limit of what may be inferrable without additional assumptions. Building on our theoretical work on the identifiability of the tree of blobs from gene quartet distributions under the Network Multispecies Coalescent model, we develop an algorithm, TINNiK, for statistically consistent tree of blobs inference. We provide examples of its application to both simulated and empirical datasets, utilizing an implementation in the `MSCquartets 2.0` R package.

**MSC Classification:** 92D15, 92D20

## 1 Background

The availability of genome-scale datasets has led to a shift in focus of methodological work in phylogenetics. The Multispecies Coalescent (MSC) model, which captures how incomplete lineage sorting (ILS) may lead to gene trees discordant with one another and a species tree, now provides the theoretical basis for many approaches to species tree inference [1–7]. However, the analysis of genomic sequence data has also made clear that using trees to model species relationships can be inadequate.

Species networks allow for the description of more complex patterns of sequence evolution produced by hybridization or other forms of lateral gene transfer. Such a network may show tree-like evolution in some parts, with other parts, called *blobs*, displaying reticulations indicating transfers of genetic material between populations.

These blobs may range in complexity from simple isolated cycles with a single reticulation to arbitrarily complex structures with numerous reticulations. Since some forms of gene transfer are believed to be more likely among closely related species, and thus occur when ILS is also present, the Network Multispecies Coalescent (NMSC) model is usually adopted to describe the combined effects of both gene transfer and ILS in the formation of gene trees [8–12].

Inference of a species network under the NMSC model, however, poses major challenges. Simultaneous inference of gene trees and species networks from sequences in a Bayesian framework is computationally demanding, with successful attempts limited to very small datasets [13, 14] of few taxa and genes. Inference of gene trees by standard phylogenetic methods, with these inferred gene trees treated as “data” for a second stage of species network inference, allows for the analysis of larger datasets. Since likelihood inference of a network still requires substantial computational effort, pseudolikelihood on summary statistics may be used instead. This approach is taken by PhyloNet [15], using gene tree rooted triples, and SNaQ [16], using gene quartets, with both requiring pre-specification of the number of reticulations. Additional speed is obtained in SNaQ by limiting networks to a level-1 structure. NANUQ, [17], also based on quartets, attains considerably greater speed by limiting statistical testing to gene tree quartets and then using combinatorial methods for network building. NANUQ also is limited to level-1 networks but can give some indication of when the level-1 hypothesis is violated. Finally, PhyNEST [18] also performs level-1 quartet-based pseudolikelihood inference, but uses genomic site pattern data with the assumption that all sequences on all gene trees were generated under the Jukes-Cantor model of site substitution.

While assuming level-1 structure is helpful computationally, as these methods show, it is unlikely to be justifiable in all biological settings. Nonetheless, some limit on network complexity is necessary for acceptable computational time, and even networks only slightly more complicated than level-1 may lack identifiability from certain data types [19].

The algorithm presented in this work takes a step toward addressing this problem, by inferring the *tree of blobs* [20] of an arbitrary species network. In this tree only cut edges of the network remain while the blobs are shrunk to nodes. (See, for example, Figure 1.) Thus multifurcations in the tree of blobs represent potentially quite 2 complicated reticulated structures for which no detailed description is given. This is similar to a “soft polytomy” in an inferred gene tree, which rather than representing a detailed evolutionary relationship indicates merely an inability to obtain the true resolution. The tree of blobs thus serves as a partial answer to how we can efficiently infer species networks, isolating those parts of the network which require additional tools to be applied — and developed — for inferring the detailed reticulate relationships.

**Fig. 1.**
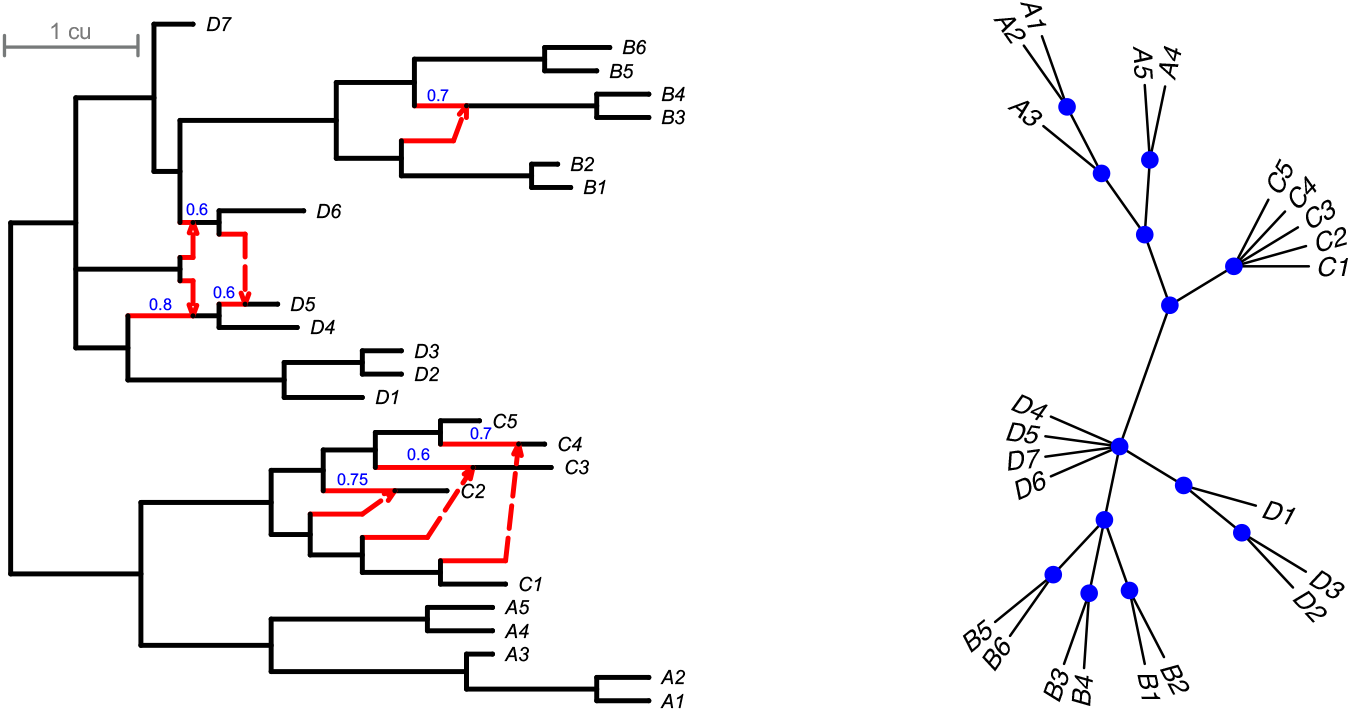
(L) A species network *𝒩* ^+^ with branch lengths in coalescent units, and (R) its tree of blobs. Hybrid edges of *𝒩* ^+^ are red, with hybridization parameters in blue above the major hybrid edges. The extended Newick string is given in Appendix A. The network *𝒩* ^+^ is non-binary, non-level-1, non-ultrametric and non-tree-child with a 7-blob, a 6-blob, and a 4-cycle. These blobs correspond to nodes of degree 7, 6, and 4 in the true tree of blobs on the right, where blobs are shown as blue dots. The network *𝒩* ^+^ is used in simulations in Section 5.1 to validate the TINNiK algorithm.

In a previous work [21], as a byproduct of proving the theoretical identifiability of the tree of blobs under the NMSC, an algorithm to infer it from gene trees was sketched. Hypothesis tests on counts of quartets displayed across gene trees for sets of 4 taxa allow the putative determination of some *blob quartets*, that is, of sets of 4 taxa that are best related by a single blob, and other quartets that could be related by a tree. Maximum likelihood then allows for assignment of a tree topology to the latter. However since not all blob quartets can be identified directly from gene data on single 4-taxon sets, a new combinatorial inference rule is needed to combine data for multiple 4-taxon sets. As was shown in [21], repeated application of this rule is sufficient to correctly identify all blob quartets. With all blob quartets known, and tree topologies assigned to other sets of 4 taxa, we use a certain intertaxon distance that can be computed from this information and which, assuming no error, exactly fits the tree of blobs. Standard distance tree building methods which are robust to some error can then be used to infer the tree of blobs from data.

We provide a detailed algorithm for a fast implementation of this method, called <u>T</u>ree of blobs <u>IN</u>ference for a <u>N</u>etwor<u>K</u>, or TINNiK,^1^ as well as an implementation in the `MSCquartets` R package, v. 2.0 [22, 23]. We further show TINNiK provides a statistically consistent estimate of a network’s tree of blobs, provided its input is a sample of gene trees under the NMSC. Since in practice the input will be gene trees inferred from sequence data, some degradation of performance is to be expected. Nonetheless, sample explorations with simulated and empirical data indicate good performance and short computation time.

We know of no other proposed algorithm for tree of blobs inference from biological data. One might consider obtaining such an estimate through a modification of the NANUQ algorithm [17], by collapsing the blobs in the NANUQ splits graph to nodes. However, while it is not hard to see this would give a statistically consistent estimator in the level-1 case, nothing is known about the theoretical behavior of doing so for more general networks.

This paper is structured as follows. Section 2 provides basic definitions and restatements of key results from [21] which underlie our algorithm. Section 3 presents a new statistical test to distinguish 4-taxon networks with a blob from those without one, with the derivation of the test distribution deferred to Appendix C. In Section 4, we present our TINNiK algorithm for the inference of a tree of blobs for a species network, and show its consistency under the NMSC model. Section 5 explores performance on both simulated and empirical datasets, with Section 6 offering concluding comments.

## 2 Networks and models

The theoretical underpinnings of the TINNiK algorithm were developed carefully in [21], so we treat the fundamental definitions and background more informally here. Readers should consult the earlier work for a more complete development.

### 2.1 Phylogenetic Networks

We denote by *𝒩*^+^ a *rooted phylogenetic network*, that is, a connected, rooted, directed graph with no directed cycles. See Figure 1 (L) for an example. Taxa in a set *X* bijectively label the leaves, the degree-1 descendants of the root. Nodes are classified as *tree* or *hybrid* according to whether exactly 1 edge or more enters them. Edges are similarly classified according to their child node. We often focus on *binary* networks, in which the root is degree 2 and all internal vertices are degree 3. For formal definitions of particular classes of networks, including level-*k*, we recommend [24].

A metric structure on the network specifies numerical parameters for the NMSC model. Edge lengths are measured in *coalescent units* (units of generations/population size), with tree edge lengths positive. Hybrid edges have non-negative lengths (with length 0 modeling instantaneous jumping of a lineage from one population to another). *Hybridization parameters* are positive probabilities that a gene lineage at a hybrid node follows a particular hybrid edge as it moves backward in time toward the root.

The *least stable ancestor* (LSA) of a network is the lowest node through which any path from the root to any taxon must pass. While a network may have a complicated structure above its LSA, our methods do not give us any information about this, nor about the location of the LSA. For this reason, our focus is on the *semidirected phylogenetic network 𝒩*^*−*^, obtained from *𝒩*^+^ by deleting nodes above the LSA, undirecting all tree edges, and suppressing the LSA if it became a degree-2 node. Note that *𝒩*^*−*^ is unrooted, but retains the directions of all hybrid edges. Provided no ambiguity results, the symbol *𝒩* may denote either *𝒩*^+^ or *𝒩*^*−*^ for simplicity.

A rooted phylogenetic network *𝒩*^+^ on a set of taxa *X* induces a network 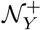 on any subset *Y⊂ X*, by retaining only edges and nodes ancestral to at least one taxon in *Y*. Induced networks on 4-taxon sets will play a particularly important role in this work.

### 2.2 Blobs

A *cut edge* in a graph is one whose deletion increases the number of connected components of the graph. The following definition also applies to general graphs.

#### Definition 1.

*A* blob *on a graph is a maximal connected subgraph with no cut edges. An edge in a graph is* incident to a blob *if exactly one of its endpoints is in the blob. A blob is an m*-blob *if it has exactly m incident cut edges*.

While blobs may have complicated structures, the simplest possible form is a single node, which is a *trivial blob*. For example, on a tree all blobs are trivial. The next simplest form a blob may have is that of an (undirected) cycle.

#### Definition 2.

*[20] The* strict tree of blobs, *𝒯* (*𝒩*), *for any connected graph, 𝒩, is the tree obtained by contracting each of the network’s blobs to a vertex, that is, by removing all of the blob’s edges and identifying all its vertices*.

A blob with *m* incident cut edges in a network leads to an *m*-multifurcation in the strict tree of blobs, so 2-blobs give degree-2 nodes. Since our methods cannot detect 2-blobs, we use a variant of the general notion of a tree of blobs.

#### Definition 3.

*The* reduced unrooted tree of blobs, *𝒯* = *𝒯*_*rd*_(*𝒩*^*−*^), *of a rooted phylogenetic network 𝒩*^+^ *is obtained from the strict tree of blobs of the semidirected network 𝒩*^*−*^ *by suppressing all degree 2 nodes*.

For the remainder of this work, the strict tree of blobs plays no role. Therefore, we refer to the reduced unrooted tree of blobs simply as the ‘tree of blobs *𝒯*.’ See Figure 1 (R) for an example.

### 2.3 Quartets

We use two distinct classifications of sets of 4 taxa as quartets, expressing different relationships of these sets to the structure of a network. The first is the standard notion of a quartet [25] in which, for instance, *ab*|*cd* refers to an unrooted topological tree with a cut edge separating the taxa *a, b* from *c, d*, and *abcd* refers to the star tree.

A different notion of quartet captures the relationship of a set of 4 taxa to the blobs of a network. A set of 4 taxa *defines* a blob *ℬ* if there are 4 disjoint undirected paths from *ℬ* to these taxa. The taxa define *ℬ* precisely when deleting *ℬ* and its incident edges leaves the 4 taxa in distinct connected components.

#### Definition 4.

*[21] A set Q* = {*a, b, c, d*} *of 4 taxa on an n-taxon network is a* Blob quartet, *or* B-quartet, *if there is a blob on the network which is defined by Q*.

*If a set of 4 taxa is not a B-quartet on a network, then it is a* tree-like quartet, *or* T-quartet.

A B-quartet *Q* = {*a, b, c, d*} on *𝒩*^+^ induces the unresolved quartet topology *abcd* on the tree of *𝒯* blobs of *𝒩*^+^, while a T-quartet induces a resolved quartet topology on *𝒯*. Note, however, that a B-quartet on *𝒩*^+^ may become a T-quartet on an induced network 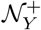. For instance, if *𝒩*^+^ is a 5-taxon network with a single blob which is a 5-cycle (i.e., a 5-*sunlet* network) and *Q* is the 4 taxa not descended from the hybrid node, then *Q* is a B-quartet on *𝒩*^+^, but a T-quartet on 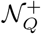. See, for example, Figure B1 in Appendix B. In contrast, T-quartets on a large network remain T-quartets on induced subnetworks.

### 2.4 Quartet Concordance Factors and the B-quartet inference rule

The NMSC model on a metric phylogenetic network determines a distribution of binary metric gene trees, and, through marginalization, distributions of binary topological gene trees on subsets of taxa. For subsets of 4 taxa, these distributions have a special name.

#### Definition 5.

*Let 𝒩* ^+^ *be a metric rooted phylogenetic network on a taxon set X, and a, b, c, d* ∈ *X distinct taxa. The* (quartet) concordance factor *CF*_*ab*|*cd*_ = *CF*_*ab*|*cd*_(*𝒩*^+^) *is the probability under the NMSC model on 𝒩*^+^ *that a gene tree displays the quartet ab cd. The* (vector quartet) concordance factor, *CF*_*abcd*_ = *CF*_*abcd*_(*𝒩*^+^) *is the ordered triple*

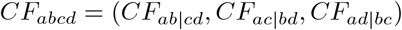

*of concordance factors of each resolved quartet on a, b, c, d*.

Since under the NMSC on any phylogenetic network all gene trees are binary and all have positive probability, the entries of *CF*_*abcd*_ for any *a, b, c, d* are positive and sum to 1.

#### Definition 6.

*CF*_*abcd*_ *is said to be* cut *if two of its entries are equal, and* strictly cut *if in addition the third entry is distinct. If CF*_*abcd*_ *is strictly cut with CF*_*ab*|*cd*_ */*= *CF*_*ac*|*bd*_ = *CF*_*ad*|*bc*_, *then we say CF*_*abcd*_ *is* strictly (*ab cd*)-cut. *If CF*_*abcd*_ *is not cut, we say it is* non-cut.

The terminology “cut” is motivated by the following theorem.

#### Theorem 1.

*[21]* (*CF-detectability of 4-blobs on 4-taxon networks*) *Consider a 4-taxon rooted binary phylogenetic network 𝒩* ^+^ *on taxa* {*a, b, c, d*} *with quartet concordance factor CF*_*abcd*_ *and tree of blobs 𝒯. Then under the NMSC model for generic parameters:*

a. *𝒯 has the quartet tree topology ab*|*cd if, and only if, CF*_*abcd*_ *is strictly* (*ab*|*cd*)*-cut*.
b. *𝒯 has the unresolved quartet topology if, and only if, CF*_*abcd*_ *is non-cut*.

In contrast to the notions of B- and T-quartets, which refer to the relationship of 4 taxa through the topology of a full network *𝒩*^+^, the notions of cut and non-cut *CF* s refer to properties of the probability distribution under the NMSC, and thus depend only on the induced 4-taxon network.

Theorem 1 shows that on 4-taxon networks there is a close correspondence between these concepts. However, on a larger network they diverge, with the following theorem giving a further tool for relating them.

#### Theorem 2.

*[21]* (*B-quartet Inference Rule*) *Consider a rooted binary phylogenetic network 𝒩*^+^ *on n taxa, n ≥* 5. *Suppose that* {*a, b, c, d*} *and* {*b, c, d, e*} *are B-quartets on 𝒩* ^+^. *If on the induced 4-taxon network any one of* {*a, b, c, e*}, {*a, b, d, e*}, *or* {*a, c, d, e*} is

a. *a T-quartet, with a, e not a cherry on the reduced unrooted tree of blobs for the induced 4-taxon network, or*
b. *a B-quartet*,

*then all of*{ *a, b, c, e*}, {*a, b, c, e*}, *and* {*a, b, c, e*} *are B-quartets on 𝒩*^+^.

The previous two theorems lead to a powerful result for application.

#### Theorem 3.

*[21] On an n-taxon rooted binary phylogenetic network 𝒩*^+^ *with generic numerical parameters, all B-quartets can be identified from the quartet CFs using CF-detectability* (*Theorem 1*) *and applications of the B-quartet Inference Rule* (*Theorem 2*).

In [21], these three theorems were the key to establishing that the tree of blobs of an arbitrary binary species network is identifiable from gene quartet concordance factors. In this work, they form the basis of an algorithm to infer that tree of blobs.

## 3 Statistical testing and estimation for cut *CF* s

A key component of the TINNiK algorithm for inference of the tree of blobs is testing gene tree data to determine which sets of four taxa are in accord with a cut *CF*. For this, we introduce a new hypothesis test.

### 3.1 Cut model testing and maximum likelihood inference

For any phylogenetic network, a *CF* is a point in the interior of the 2-dimensional probability simplex, Δ^2^ ={(*p*_1_, *p*_2_, *p*_3_) |*p*_*i*_ *>* 0, *∑p*_*i*_ = 1}

#### Definition 7.

*The* cut model *comprises those points in* Δ^2^ *representing cut CF s, that is* {(*p*_1_, *p*_2_, *p*_3_) ∈ Δ^2^ |*p*_*i*_ = *p*_*j*_ *for some i* ≠ *j*}, *as depicted in Figure 2* (*L*).

**Fig. 2.**
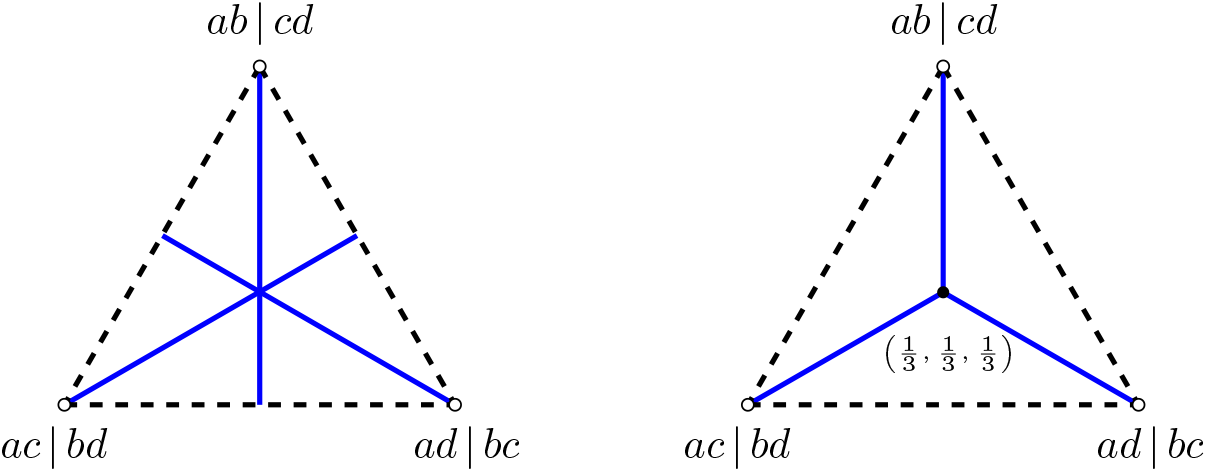
Geometric view of *CF* s for 4-taxon network models, with dashed lines outlining the simplex Δ^2^. Each point in Δ^2^ arises as a *CF* under the NMSC, even when restricting to level-1 networks [26]. (L) The cut model consists of 3 blue line segments, with each formed by *CF* s arising from 4-networks with a specific resolved tree of blobs topology. *CF* s off the cut model arise only from networks with unresolved trees of blobs. (R) The *T* 3 submodel is those cut *CF* s with smallest entry occurring exactly twice. Using it as the null hypothesis in the TINNiK algorithm may lead to more sets of 4 taxa initially judged as B-quartets than using the cut test.

**Fig. 3.**
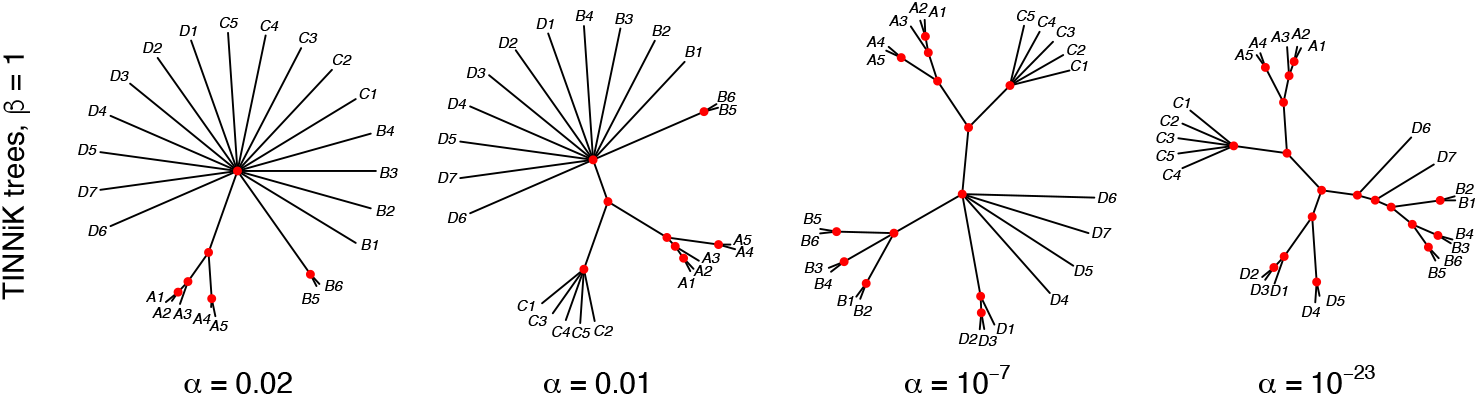
Four TINNiK trees of blobs for a simulated sample of *n* = 1000 gene trees on network *𝒩* ^+^ of Fig. 1 (L) with *k* = 1 (moderate ILS), for *β* = 1 and *α* = 0.02, 0.01, 10^*−*7^, 10^*−*23^. Increasing resolution as *α* is decreased is typical. The true tree of blobs (third from left) is inferred for a large range of test levels *α* ∈ [10^*−*7^, 0.001]. Although the rightmost tree is over-resolved, each split is compatible with a tree displayed on *𝒩* ^+^.

**Fig. 4.**
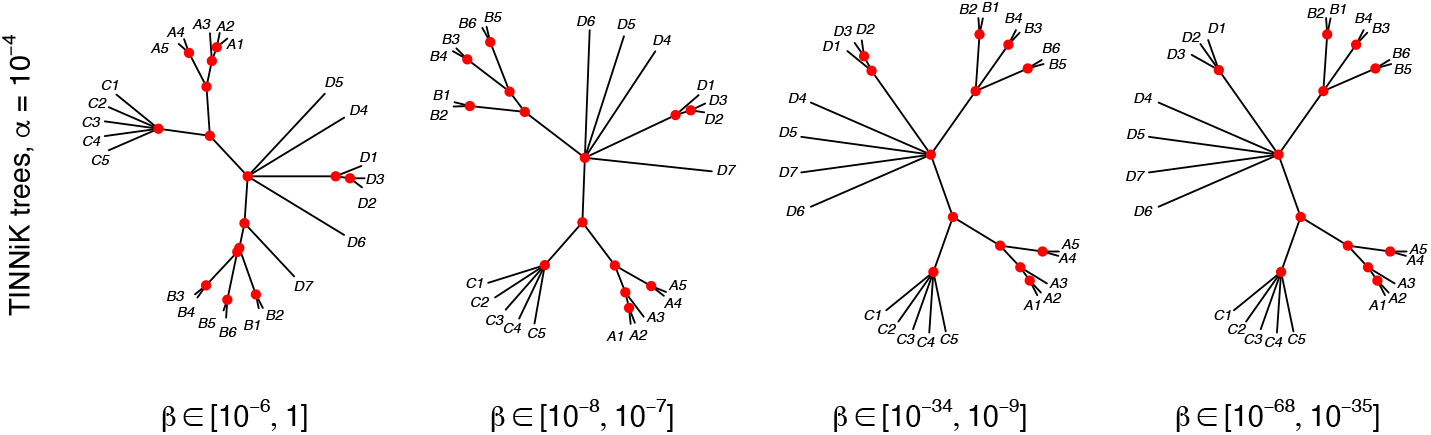
The TINNiK tree of blobs for fixed *α* = 10^*−*4^ and various *β* for a simulated sample of *n* = 1000 gene trees for *𝒩* ^+^ with *k* = 0.5 (high ILS). Decreasing *β* results in less resolution in the tree of blobs. From left to right: only the *A*- and *C*-groups are correct; all blobs except the *B*-blob are correct; the TINNiK tree of blobs is correct; the *D*-group lacks sufficient resolution. For even smaller *β* the TINNiK tree of blobs degrades to a star tree.

Data relevant to a *CF* is collected in the form of a *quartet count concordance factor* (*qcCF*) [27], a vector of counts

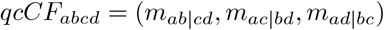

of the three resolved unrooted topological quartet gene trees, which for the taxon set {*a, b, c, d*}are assumed to be independently drawn from the NMSC. In practice, these could be quartet trees individually inferred from sequence data for different genes, or quartet trees displayed on inferred gene trees on more taxa. However, our development of a statistical test assumes no inference error is present.

With total sample size *m* = *m*_*ab*|*cd*_ + *m*_*ac*|*bd*_ + *m*_*ad*|*bc*_, the *empirical concordance factor*, which consistently estimates the concordance factor, is

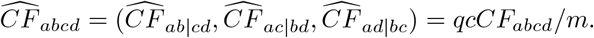

Viewing 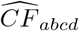 as a point in the simplex, closeness to the cut model lines lends informal support that the true *CF*_*abcd*_ is cut, while a greater distance supports that *CF*_*abcd*_ is non-cut. For judging closeness, however, one must take the sample size *m* into account.

To formulate a formal hypothesis test, fix four taxa *a, b, c, d*, and the data *qcCF*_*abcd*_ = (*m*_*ab*|*cd*_, *m*_*ac*|*bd*_, *m*_*ad*|*bc*_). Assuming *qcCF*_*abcd*_ arises as a trinomial sample from the distribution specified by some true *CF*_*abcd*_, consider null and alternative hypotheses:

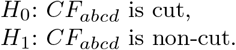

For a test statistic, we use the likelihood ratio statistic for the null and alternative models, with Appendix C.1 presenting the necessary calculations for the test.

Because the cut model has a singularity at (1*/*3, 1*/*3, 1*/*3) (Figure 2 (L)), standard assumptions underlying the routine use of the 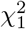 distribution for judging the test statistic are violated there. However, *CF* points near the centroid include those for trees and networks with short internal branches, and thus include some of those of the greatest interest to researchers. Building on work in [28], we thus develop an alternative testing distribution that takes into account this geometry of the cut model. Appendix C.2 presents its derivation and Appendix C.3 simulations illustrating its improved performance over the 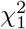 distribution near and at the cut model singularity.

### 3.2 The *T* 3 model and testing

An *anomalous quartet* {*a, b, c, d*}is one whose *CF* is *ab*|*cd* cut of the form (*p, q, q*) with *q ≥* 1*/*3. Graphically, this means the *CF* lies on the cut model depicted in Figure 2 (L), but not on the *T* 3 model shown in Figure 2 (R). In [17], anomalous quartets for level-1 networks were investigated and shown to require a 3-cycle with two taxa descended from the hybrid node, and somewhat extreme numerical parameters which seem unlikely biologically. Further investigation by Ané et al. [29] suggests that anomalous quartets for more complex networks are also not likely to be common. For this reason, when inferring a tree of blobs it can be reasonable to assume that an unknown network has no anomalous quartets and use a *T* 3 hypothesis test for *CF* s, rather than a cut test. The *T* 3 test, developed in [28], is a backbone for the NANUQ method [17] for inferring level-1 topological networks under the NMSC. Using the *T* 3 test one might infer more B-quartets than using the cut test, as any 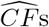 near the cut model line segments of the form (*p, q, q*) with *q >* 1*/*3 might support the null hypothesis for the cut test, but be rejected by the *T* 3 test and flagged as indicating 4-blobs. Thus, using the *T* 3 test can produce a less resolved tree of blobs than the cut test. While this is not a conservative approach in the sense of hypothesis testing (since it may lead to more rejections of a null hypothesis of a tree-like quartet relationship), the inferred tree of blobs it produces is a more cautious one that possibly avoids depicting erroneous resolution.

Although our implementation of TINNiK in `MSCquartets` has a default option of the *T* 3 test, we recommend performing data analysis with both tests. For many datasets we have found they give identical results, since they either infer the same initial B-quartets, or the B-quartet inference rule compensates for missing some of these initially using the cut test. When the results of using the two tests differ, investigating why that occurred may provide more insight into the data.

## 4 The TINNiK algorithm for inference of the tree of blobs

Building on the theorems and hypothesis tests from previous sections, we present a detailed algorithm, <u>T</u>ree of blobs <u>IN</u>ference for a Species <u>N</u>etwor<u>K</u>, or TINNiK, for inferring the reduced unrooted topological tree of blobs of a network *𝒩*^+^ from multigene data. We then analyze its running time and show its statistical consistency under the NMSC model for binary *𝒩*^+^.

TINNiK applies hypothesis tests to classify empirical *CF* s as cut or non-cut, giving quartet trees of blobs, using Theorem 1. Then it repeatedly but efficiently uses the inference rule of Theorem 2 to infer all B-quartets for a network. TINNiK’s next step is to use a quartet-based intertaxon distance formula [6] to convert B- and T-quartet information to a distance approximately fitting the topological tree of blobs. Then an inferred tree of blobs can be obtained by any of a number of well-known tree-building algorithms such as Neighbor-Joining [30], DescentTree [31], or FastME [32]. If the quality of the input data is unknown, or its fit to the NMSC model is doubted, we recommend the use of the Neighbor-Net algorithm [33] to confirm the distance reflects a strong tree signal before tree building.

One concern for algorithm design is how to handle empirical *CF* s that are near (1*/*3, 1*/*3, 1*/*3). These might arise from either a true multifurcation in a network (a hard polytomy), a “near multifurcation” of a resolved subnetwork with short internal edges (a soft polytomy), or from complex blobs with longer edges. TINNiK treats all *CF* s judged by a “star tree” hypothesis test to be close to (1*/*3, 1*/*3, 1*/*3) as B-quartets. While this is a natural approach, it does mean that the inferred tree of blob’s structure may reflect both true blobs and further multifurcations due to data quality that is insufficient to resolve some cut edges. This may cause inferred blobs to be larger than true ones, but only when the data is inadequate to obtain greater resolution.

### 4.1 Algorithms

The B-quartet Inference algorithm takes as input a table of quartet count concordance factors (*qcCF* s). Working from a collection of *m* gene trees, all on the full set *X* of *N* taxa, this quartet table can be produced in time *𝒪* (*mN* ^4^). Although under the NMSC model all gene trees are fully resolved, in applications some inferred trees may not be, but these can be handled either by discarding their unresolved quartets or assigning uniform “counts” of 1/3 to each resolved topology, as discussed in [27].

In order to access entries of this table rapidly, without scanning it in its entirety, we require that its rows, corresponding to sets of 4 taxa, be ordered so that the index for any set can be computed directly and quickly. We choose to order the sets of four taxa by *lex order*. In more detail, this means that if the taxa are designated by the numbers 1, 2, 3 …, *N*, and a set of 4 taxa is designated by a “word” of the four numbers in ascending order, then these words are ordered lexicographically using the usual order on natural numbers. Thus the first few sets are ordered as

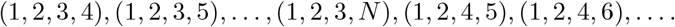

In lex order, the index for a particular set of 4 taxa is given by the formula (Corollary 3.22, [34])

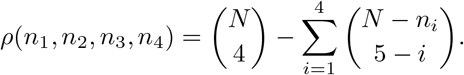

The computational simplicity of this formula allows for its rapid evaluation. Tabulating all binomial coefficients that might be needed in the formula in advance, so they are computed only once, requires time *𝒪* (*N*). After this, however, the index for any set of 4 taxa can be computed in time *𝒪* (1).

Two hypothesis tests are used in the algorithm. First, we use a star tree test to determine whether each *qcCF* is consistent with a 4-polytomy for each induced 4-taxon network. The null hypothesis is that the *CF* for the 4 taxa is (1*/*3, 1*/*3, 1*/*3), with the alternative its complement in the simplex. A standard 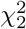 test at some level *β* on the likelihood ratio statistic is performed. Failure to reject the null suggests either a true 4-polytomy, or lack of sufficient information to infer a resolution. In addition, one of the cut or *T* 3 hypothesis tests described in Section 3 is used in the algorithm to decide when a vector *qcCF* for four taxa is in accord with a cut relationship. More formally, rejecting the null hypothesis of this test at level *α* is interpreted as indicating a non-cut *CF*. By Theorem 1, this is evidence the 4 taxa form a B-quartet on the induced 4-taxon species network.

The following algorithm applies these tests and the inference rule for B-quartets of Theorem 2.

#### Algorithm.

(*B-quartet Inference*)

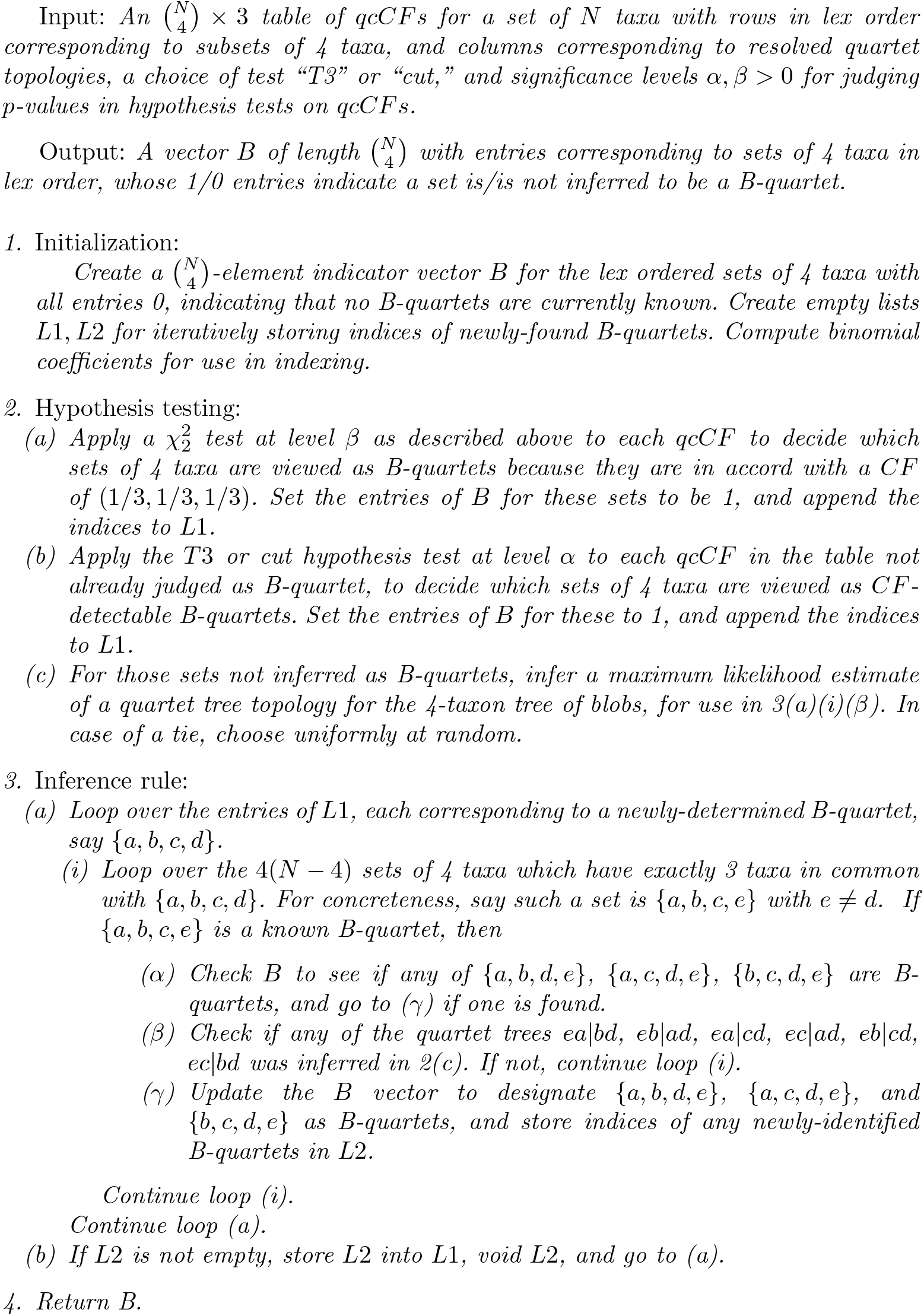

In this algorithm, step 2 implements the theoretical *CF*-detectability result of Theorem 1, while step 3 implements the B-quartet inference rule of Theorem 2. The algorithm eventually considers every pair of B-quartets sharing three taxa, since any time a new one is discovered it is compared to all quartets that share three taxa with it. If one of these is a not-yet-inferred B-quartet, then this pair will be compared again later, once that quartet is inferred as a B-quartet. Theorem 3 therefore ensures the looping of step 3 can determine all B-quartets, assuming sufficient data in accord with the NMSC.

Steps 1 and 2 can each be accomplished in time *𝒪* (*N* ^4^). Since there can be at most 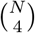 B-quartets that can appear in the lists *L* through all passes through step 3a, and each is compared in step 3(a)i to *𝒪* (*N*) other sets of 4-taxa, all applications of step 3 require time at most *𝒪* (*N* ^5^). Thus the total time complexity is *𝒪* (*N* ^5^).

To estimate the tree of blobs for a network, we seek a tree that displays unresolved quartet trees for all B-quartets and a resolved quartet tree with the topology estimated by maximum likelihood for all T-quartets. Note that the estimate of the topology is recorded when step 2c of the B-inference algorithm is performed, so we treat this as known.

Estimating the tree of blobs is now an instance of a supertree problem, with input all trees on 4 taxa, with the trees for B-quartets unresolved. To address this, we take the approach introduced in [6], in which an intertaxon distance is defined using quartet data – including that for unresolved quartets – to compute an intertaxon distance. Assuming perfect information, this distance would exactly fit the unknown tree. While inference may well lead to some incorrect quartets, distance-based tree construction methods that behave well under some noise can be used to return an inferred tree of blobs. Because of possible error in some quartets, and hence in the computed quartet distance, the inferred tree may not show exact polytomies, but rather some resolutions of them with short edges. It may thus be desirable to reduce to zero all edge lengths smaller than some cutoff *δ*. Theory behind such a cutoff will be discussed in the next subsection, in the proof of Theorem 4.

The full TINNiK algorithm we now outline takes as input a collection of gene trees on the taxa *X*, and returns an inferred topological tree of blobs for the network parameter which under the NMSC model produced those gene trees.

#### Algorithm.

(*TINNiK*)

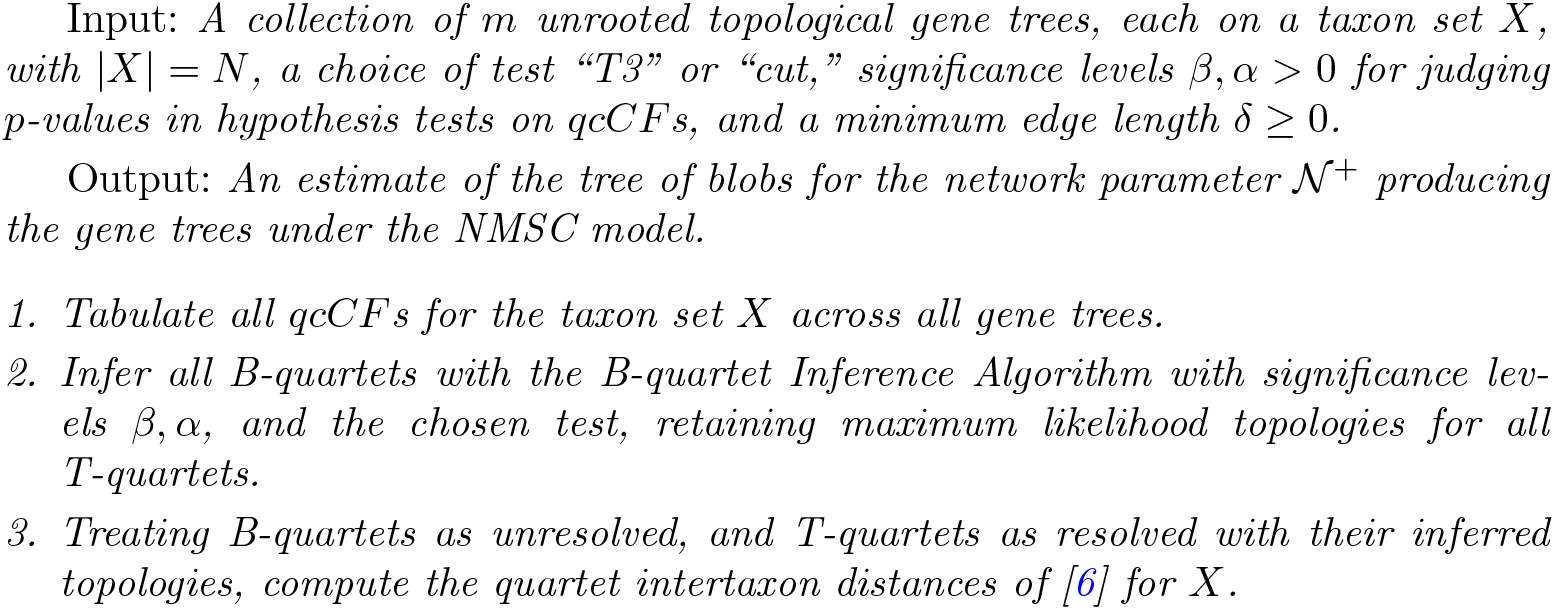
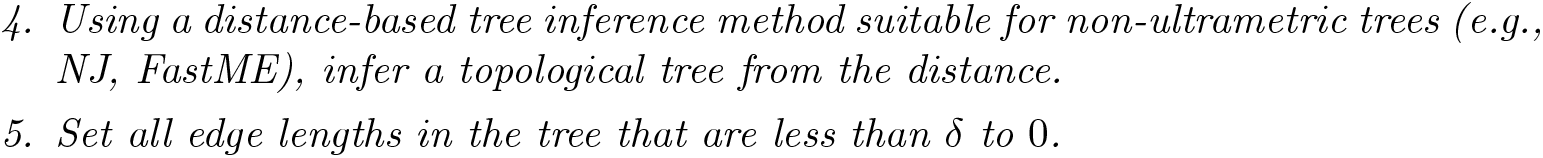

The computational times for steps 1-5 using NJ in step 4 are, respectively, *𝒪* (*mN*_*4*_), *𝒪* (*N*_5_), *𝒪* (*N*_4_), *𝒪* (*N*_3_), *𝒪* (*N*), for a combined *𝒪* ((*m + N*)*N*_4_). We report computational times in practice in Table 1, when analyzing TINNiK on simulated and real data.

**Table 1.**
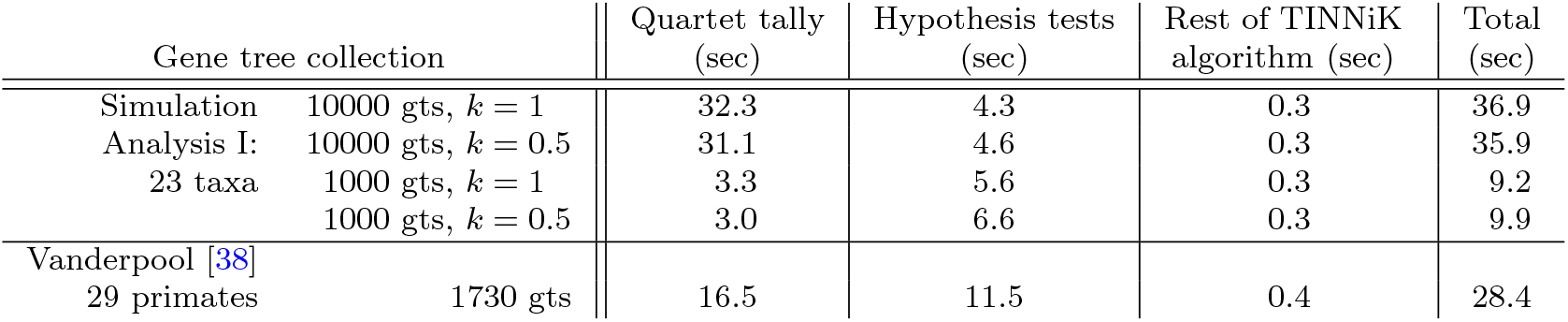
Runtimes (averaged over ten runs) for the TINNiK algorithm in MSCquartets, on a 2020 Macbook 2 GHz Quad-Core i5, 32 GB RAM. Test levels are *α* = 0.001 for the *T* 3 test, and *β* = 0.95.

| Gene tree collection |  | Quartet tally<br>(sec) | Hypothesis tests<br>(sec) | Rest of TINNiK<br>algorithm (sec) | Total<br>(sec) |
| --- | --- | --- | --- | --- | --- |
| Simulation | 10000 gts, $k = 1$ | 32.3 | 4.3 | 0.3 | 36.9 |
| Analysis I: | 10000 gts, $k = 0.5$ | 31.1 | 4.6 | 0.3 | 35.9 |
| 23 taxa | 1000 gts, $k = 1$ | 3.3 | 5.6 | 0.3 | 9.2 |
| | 1000 gts, $k = 0.5$ | 3.0 | 6.6 | 0.3 | 9.9 |
| Vanderpool [38] |  |  |  |  |  |
| 29 primates | 1730 gts | 16.5 | 11.5 | 0.4 | 28.4 |

The TINNiK algorithm can also be applied when gene trees have missing taxa, provided each subset of 4 taxa occurs on at least one gene tree, so the qcCF is not the zero vector.

### 4.2 Statistical consistency

It is desirable that inference algorithms produce statistically consistent estimators. In this context, informally this means that given data (*m* gene trees) produced under the NMSC model on a species network, the probability of obtaining the correct tree of blobs approaches 1 as the amount of data approaches infinity. However, since the algorithms assume generic numerical parameters, and there are several other algorithm inputs, *α, β, δ*, a precise statement of an appropriate notion of consistency is more complicated. We proceed similarly to how consistency was addressed for the quartet-based NANUQ algorithm for inferring a level-1 species network in [17]. For simplicity, we also restrict to the case that the data is *m* gene trees, each on the full taxon set *X*, since generalizing from this is straightforward.

Before stating our formal consistency theorem, we describe explicitly what we mean by generic parameters. For a fixed topological binary species network, it is possible that the *CF* for an induced 4-taxon network with a 4-blob may be a cut *CF*. By Theorem 1, however, for each such topological 4-network the *CF* is non-cut for all parameters except those in a measure-0 subset of its numerical parameter space. Since for any full network there are only finitely many induced topological 4-networks and the finite union of measure-0 sets has measure zero, for generic parameters (*i*.*e*., those outside this measure-0 set) on the full network, all *CF* s for quartets inducing 4-blob networks will be non-cut.

We also need that for generic parameters the *CF* of an induced 4-taxon network is not (1*/*3, 1*/*3, 1*/*3). In the case of a 4-blob, this follows from the last paragraph. But since a 4-network without a 4-blob may have a *CF* with equal entries (e.g., a 3_2_-cycle network [26]), more argument is needed that this does not occur generically. Note that if all hybridization parameters on a binary 4-network *𝒬* without a 4-blob are 0 or 1, then *𝒬* is essentially a resolved tree, for which *CF* ≠ (1*/*3, 1*/*3, 1*/*3). Analyticity of the parameterization then implies this inequality for generic parameters. Mimicking the argument above shows that no induced 4-taxon network has *CF* = (1*/*3, 1*/*3, 1*/*3) for generic parameters on the full binary network.

These preliminary observations are used to prove the following:

#### Theorem 4.

*For generic numerical parameters on a binary phylogenetic network 𝒩*^+^, *the TINNiK Algorithm using the cut test provides a statistically consistent estimate of the topological tree of blobs 𝒯* = *𝒯*_*rd*_(*𝒩*^*−*^) *under the NMSC. Specifically, there exists a sequence α*_*m*_*→*0 *such that for any β >* 0 *and* 2 *> δ ≥* 0, *the TINNiK algorithm on a set of m gene trees independently drawn from the NMSC model on a binary species network 𝒩* ^+^ *will, with probability →* 1 *as m → ∞, infer 𝒯*.

*Proof*. We restrict to generic parameters ensuring that all induced 4-networks with 4-blobs have non-cut *CF* s, and no induced 4-taxon network has *CF* = (1*/*3, 1*/*3, 1*/*3). First consider step 2a of the B-quartet Inference algorithm. With generic parameters, for each set of 4 taxa the probability the 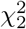 test with significance level *β* will reject the null hypothesis that a *CF* is (1*/*3, 1*/*3, 1*/*3) approaches 1 as *m→ ∞* Since there are only finitely many 4-taxon subsets, the probability goes to 1 that this null hypothesis will be rejected for all. This holds regardless of the chosen value of *β >* 0. In step 2b of the B-quartet Inference algorithm, the role of *α* in the cut test is more subtle, since if it is held fixed then we expect to erroneously reject the null model in a fraction *α* of all applications for each set of 4 taxa. To make such false negatives less common, we consider sequences of levels *α*_*m*_*→* 0 as the number of gene trees *m→ ∞* The likelihood ratio statistic is judged using the distribution of Propositions 5 and 6 of Appendix C.2. If a true *CF* is cut, then as *m→ ∞*, the parameter *µ*_0_ of that distribution goes to and the distribution converges to the 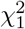. This holds even using the MLE in place of the true parameter. To ensure that the probability of failing to reject the null hypothesis approaches 1 as *m → ∞*, it is enough to choose any sequence of significance levels with *α*_*m*_ *→* 0.

In contrast, if a true *CF* = (*p*_1_, *p*_2_, *p*_3_), is non-cut and hence not in the null model, let (*m*_1_, *m*_2_, *m*_3_) denote a *qcCF* under the NMSC with sample size *m* = *m*_1_ +*m*_2_ +*m*_3_, and 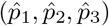 the MLE of the *CF* under the null model. Without loss of generality, assume the MLE is on the vertical line segment of Figure 2 (L), so that

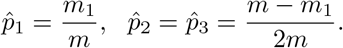

Using the formulas of Appendix C.1, the likelihood ratio statistic is then *λ* = *λ*_*m*_ = *−*2[*m*_1_ log *m*_1_ + (*m − m*_1_) log((*m − m*_1_)*/*2)

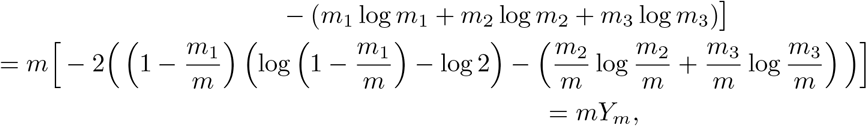

where the random variable *Y*_*m*_ converges in probability to

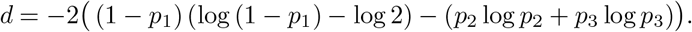

Moreover, *d >* 0 since the unconstrained likelihood has a unique maximum at (*p*_1_, *p*_2_, *p*_3_).

Now for any *η >* 0 there exists an *M* such that for *m > M*, ℙ(*Y*_*m*_ *> d/*2) *>* 1 *− η* and thus that ℙ(*mY*_*m*_ *> md/*2) *>* 1 *− η*. Since *η* was arbitrary, as *m → ∞*, ℙ(*λ*_*m*_ *> md/*2) *→* 1.

Let 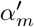 be the probability that 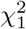-distributed random variable is greater than *md/*2, so 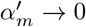 Then since the test distribution converges to the 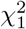, the probability of rejecting the null hypothesis is greater than 1*− η* for sufficiently large *m*. Thus the probability of rejecting the null approaches 1.

While the value of *d* depended upon the particular 4-taxon set under consideration, since there are only finitely many such sets, by choosing *α*_*m*_ as the maximum of the 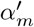 we obtain a sequence of significance levels that with probability approaching 1 as *m→ ∞* ensures the cut test will reject the null model for all induced 4-networks with a 4-blob and fail to reject it for all others. The argument so far has shown that with our choice of the *α*_*m*_ the hypothesis tests will lead us to correctly conclude that true *CF* s are cut or non-cut, with probability approaching 1 as *m→ ∞*.

When the true *CF* for 4 taxa is cut, its value is consistently inferred by maximum likelihood, and thus the topology of the 4-taxon reduced unrooted trees of blobs is as well. Thus as *m→ ∞*, with probability approaching 1, the remaining deterministic steps of the B-quartet Inference algorithm then correctly infer all B-quartets.

From this information on B-quartets and T-quartet topologies, TINNiK computes an intertaxon distance exactly fitting the network’s tree of blobs. With no error in the distances, NJ or other tree-building algorithms recover the tree exactly.

Note that *δ* played no role in this argument so far, since its purpose in the algorithm is to suppress some error which, with probability approaching 1 as *m→ ∞* is not present. We must however verify that *δ* has no detrimental effects in this asymptotic result. Reviewing [6], one sees that internal branches of a tree endowed with the quartet distance always have length of at least 2. Thus any 0*≤δ <* 2 will have no effect when all B- and T-quartets are properly determined.

### 4.3 TINNiK test levels and graphical output

When TINNiK’s hypothesis tests are applied, many sets of 4 taxa will overlap, so the *CF* s are not independent. Although a Bonferroni correction for multiple tests can be applied, controlling the family-wise error rate, we do not do so, as this is always equivalent to choosing a smaller significance level. Indeed, when the method is applied to inferred gene trees, which have unknown error, a fully justifiable formal correction is not known.

However, when a TINNiK analysis is reported for an empirical dataset, it should always include the values of *α, β* used, and whether the “cut” or “T3” test was used. Ideally, the gene trees that were used should be made publicly available, for reproducibility, as gene trees inferred by different methods might produce a different tree of blobs.

In regard to graphical output, the tree of blobs could be drawn in the usual way for phylogenies, with nodes rendered as points, but we recommend a modification. Depicting each internal node as a disk or ball gives visual emphasis that the nodes represent blobs with potentially complicated structures. Even degree-3 nodes should be shown this way, since non-node 3-blobs may exist. The implementation of TiNNiK in `MSCquartets` follows this graphical style, using red disks on an inferred tree of blobs. Finally, although any planar drawing of the tree of blobs necessarily orders the edges emanating from a blob in some way, this circular order is essentially arbitrary.

The true network may not even be embeddable in the plane without crossings, in which case no unique order is even determined. Although for certain networks (level-1, or, more generally, outer-labelled planar [19]) a unique circular order exists, TINNiK does not seek to find it, much less impose it on the tree of blobs. Viewers of a tree of blobs should keep this in mind when seeking biological insight.

## 5 Simulations and Applications

We present analyses of both simulated and empirical gene tree data, using the implementation of the TINNiK algorithm in the `MSCquartets 2.0 R` package [22, 23]. Its primary functions, TINNIK and TINNIKdist, utilize C++ code with the Rcpp package [35] for increased speed.

Datasets of gene trees were simulated under the NMSC on various networks using `PhyloCoalSimulations` [36]. As true samples under the NMSC, these do not have the gene tree inference error expected in empirical analyses. For analyses of empirical datasets, we used gene trees inferred and made publicly available by the researchers who originally analyzed them.

### 5.1 Simulations

A first set of simulations, analyzed in Sections 5.2.2 and 5.2.3, uses the model network *𝒩*^+^ of Figure 1. This network on 23 taxa with 7 hybrid nodes has some complicated features (*e*.*g*., non-binary, non-tree-child [24]), with a tree-like cluster (*A* taxa), and three blobs (*B*s, *C*s, *D*s). The *B*-blob is descended from the *D*-blob, while the *C*- and *D*-blobs include more than one instance of gene flow.

Gene tree samples of size *n* = 300, 500, 1000, 10000 were produced, with branch lengths scaled by factors *k* = 0.5, 1.0, 2.0, for a total of 12 simulation parameter settings. These include cases where sampling error may be significant (*n* = 300), and when short branch lengths and the resulting high ILS (*k* = 0.5) may confound reticulation signal. The largest value of *n* should approximate asymptotic behavior. We adopt the terms ‘high,’ ‘moderate,’ and ‘low’ ILS for the scaling factors *k* = 0.5, 1.0, 2.0, respectively, as a convenience. Since non-matching gene tree quartets under the NMSC on a species tree with internal branch length 1 occur with probability approximately 0.25, our ‘moderate ILS’ is arguably ‘moderately high.’ Simulated gene tree datasets were analyzed using the TINNIK function with the default *T* 3 test and varied values of *α* and *β*.

Since the *T* 3 test and the star tree test have different foci (hybridization vs. lack of resolution), in Sections 5.2.2 and 5.2.3 we investigate the effect of each test individually. For a general overview, Table 2 presents a summary of test levels *α* and simulation results for all parameter choices under the *T* 3 test, with *β* = 1 fixed (so all network quartets are treated as resolved), and illustrates the effect that small sample size and/or high ILS may have.

**Table 2.** Blob detection using TINNiK for simulated data for *𝒩* ^+^ of Fig. 1 (L). Entries give ranges for *α* on which the full tree of blobs, and individual blobs, are correctly inferred with the *T* 3 test. Interval endpoints are approximate, with dashes indicating the correct multifurcation is never inferred. For all analyses, *β* = 1.

| Number $n$<br>of gene trees | Range of $\alpha$ values, for blob detection. ($\beta = 1$) | | | |
| --- | --- | --- | --- | --- |
| | Tree of Blobs<br>fully correct | $B$-blob detected | $C$-blob detected | $D$-blob detected |
| Low ILS: $k = 2$. | | | | |
| 10000 | $[10^{-170}, 0.01]$ | $[10^{-170}, 0.01]$ | $[10^{-300}, 0.01]$ | $[10^{-170}, 0.01]$ |
| 1000 | $[10^{-15}, 10^{-4}]$ | $[10^{-16}, 10^{-4}]$ | $[10^{-49}, 0.01]$ | $[10^{-15}, 10^{-4}]$ |
| 500 | $[10^{-6}, 0.001]$ | $[10^{-7}, 0.001]$ | $[10^{-25}, 0.01]$ | $[10^{-6}, 0.001]$ |
| 300 | - | $[10^{-6}, 10^{-5}]$ | $[10^{-17}, 0.01]$ | - |
| Moderate ILS: $k = 1$. | | | | |
| 10000 | $[10^{-55}, 10^{-4}]$ | $[10^{-55}, 10^{-4}]$ | $[10^{-216}, 10^{-4}]$ | $[10^{-55}, 10^{-4}]$ |
| 1000 | $[10^{-7}, 0.001]$ | $[10^{-7}, 0.001]$ | $[10^{-23}, 0.01]$ | $[10^{-7}, 0.001]$ |
| 500 | $[10^{-4}, 0.005]$ | $[10^{-4}, 0.008]$ | $[1 \times 10^{-13}, 0.01]$ | $[10^{-4}, 0.005]$ |
| 300 | - | - | $[10^{-7}, 0.003]$ | - |
| High ILS: $k = 0.5$. | | | | |
| 10000 | $[10^{-12}, 0.001]$ | $[10^{-12}, 0.001]$ | $[10^{-88}, 0.001]$ | $[10^{-17}, 0.001]$ |
| 1000 | - | - | $[10^{-8}, 0.001]$ | - |
| 500 | - | - | $[10^{-4}, 0.001]$ | - |
| 300 | - | - | - | - |

To understand the effect of blob complexity, analyses in Section 5.2.4 use a second set of simulations on the networks of Figure 6. Network 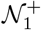 has 10 taxa and a single 7-cycle, while network 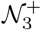 is obtained from 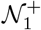 by the addition of two hybrid edges cutting across the cycle, changing the blob from level-1 to level-3. Samples of size *n* = 1000 gene trees were simulated.

**Fig. 5.**
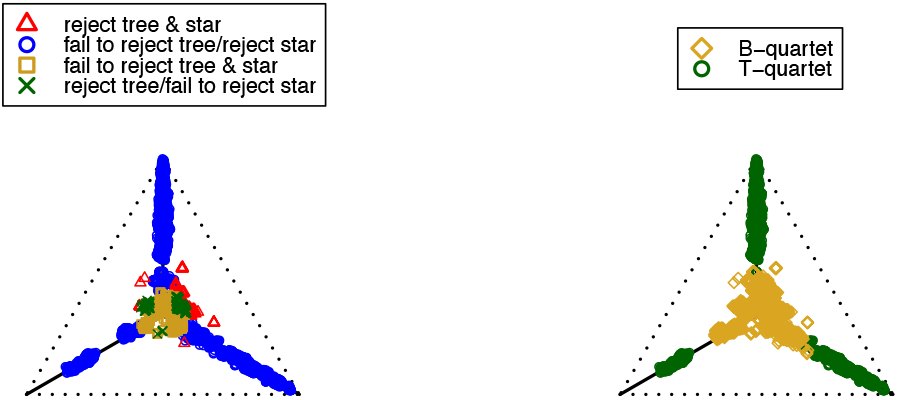
Simplex plots showing the results of hypothesis tests for *α* = 10^*−*4^, *β* = 10^*−*10^ (L) and after the application of the Inference Rule (R). B-quartet simplex plots for any *β* ∈ [10^*−*31^, 10^*−*9^] are identical, although the hypothesis test results differ.

**Fig. 6.**
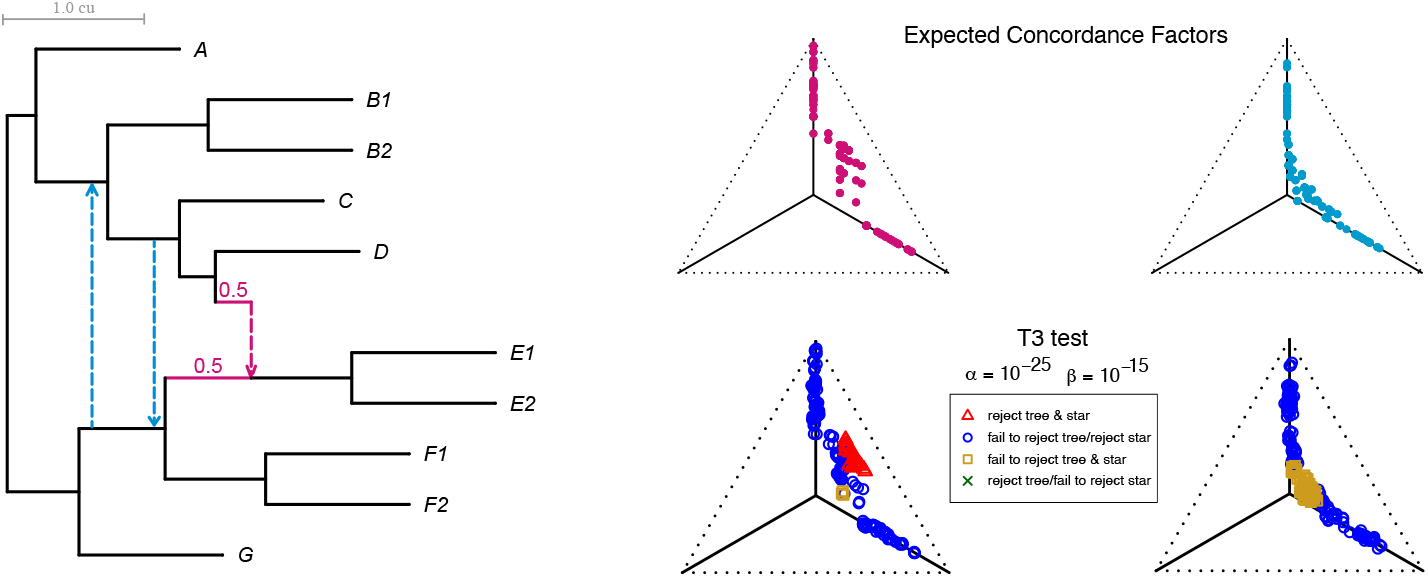
(L) Model networks in analysis III. *N*_1_ consists of black and magenta edges, and *𝒩*_3_ all edges. All hybridization parameters are *γ* = 0.5, with Newick notation given in Appendix A. (R, top) Expected *CF* s for *𝒩*_1_ (magenta) and *𝒩*_3_ (light blue). (R, bottom) Typical simplex plots for *𝒩*_1_ (left) and *𝒩*_3_ (right) displaying hypothesis test results for *α* = 10^*−*25^, *β* = 10^*−*15^. For these levels, TINNiK infers the true tree of blobs of both *N*_1_ and *𝒩*_3_, but with different initial lists of B-quartets. For *𝒩*_1_ most initial B-quartets are detected from non tree-like signal, but for *𝒩*_3_ from star-like.

A final simulation, in Section 5.2.5, generated a sample of size *n* = 10, 000 for the level-2 (2 overlapping cycles) network *𝒩*^+^ of Figure 7. Analyses were done with TINNiK and also SNaQ, which infers a level-1 network under the NMSC using pseudolikelihood on empirical quartet *CF* s [37]. SNaQ searches were done starting at the four level-1 networks displayed on *𝒩* ^+^ (obtained by deleting exactly one hybrid edge), with the user-defined maximum number of hybridizations, *h*_*max*_, set to 1 and 2. Since theory justifies SNaQ’s use only for level-1 networks, this modeling scenario violates its main assumption of network complexity, and it should not be expected to perform well. Our goal is not to point out any weakness of SNaQ, but to illustrate that TINNiK might help empiricists evaluate if an assumption made by another method is violated by contrasting its results to output from that method.

**Fig. 7.**
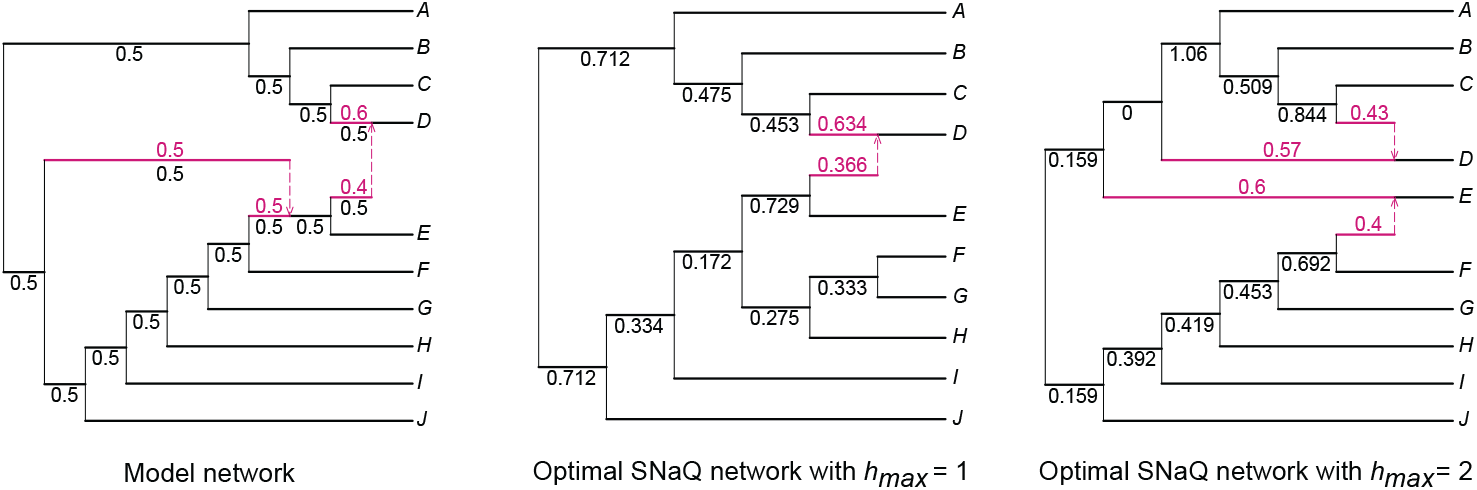
(L) A level-2 model network with star tree of blobs. Hybrid edges and hybridization parameters are in magenta, with Newick notation given in Appendix A. (C) The optimal inferred network from SNaQ with the maximal number of hybridizations constrained to *hmax* = 1, and (R) with *hmax* = 2. In both (C,R), terminal and hybrid branch lengths are absent since they are not identifiable under the NMSC from *CF* s when only a single lineage is sampled from the descendant population, and therefore are not inferred by SNaQ.

In accordance with Theorem 4, branch lengths in the TINNiK tree of blobs shorter than 2 were collapsed to zero in all analyses.

### 5.2 Results

We caution TINNiK users that one can rarely simply choose test levels *α, β* ∈ [0, 1] in advance (e.g., at the common level of 0.05) and obtain a strong analysis. Rather, a range of significance levels should be considered, in conjunction with viewing the resulting hypothesis test simplex plots and weighing one’s understanding of the extent of noise present in inferred gene trees.

Varying *α* from small to large increases the number of *CF* s interpreted as signaling hybridization, potentially causing TINNiK’s inferred tree of blobs to gain more or larger multifurcations. This can indicate which multifurcations have the strongest support. Even in simulated gene tree data, which has no model misspecification, the level of support can vary with network features such as blob complexity, hybridization parameter values, and location of a blob within the network. Simplex plots of test results, as discussed in [27], can help users choose values of *α* that give good separation of plotted *CF* s into tree-like and non-tree-like clusters.

Varying *β* from small to large decreases the number of *CF* s interpreted as indicating a star tree, potentially causing the inferred tree of blobs to be more resolved. If few *CF* s are plotted near the centroid (1/3,1/3,1/3) of the simplex, the value of *β* has little impact over a wide range. However, if many *CF* s are near the centroid, *β*’s value can be quite impactful. In some empirical datasets that have been studied for signs of hybridization we have found *CF* s so tightly clustered near the centroid that whether any signal for hybridization exceeds likely gene tree inference error seems debatable. Again, simplex plots of test results for various *β* values can be a helpful guide.

#### 5.2.1 Empirical runtimes

Representative runtimes are shown in Table 1. These were found using gene trees as input, and do not include the time to infer trees from sequence data. Runtimes for other *α, β* are similar, although the tallying of quartets need only be done once.

All times shown are a matter of seconds. In particular, TINNiK is much faster than SNaQ or PhyNEST which infer level-1 networks, and PhyloNet which seeks an arbitrary network. TINNiK’s runtime is on par with NANUQ’s for inferring a level-1 network topology. However, TINNiK gives meaningful (though coarse) output quickly without assumption on network level.

#### 5.2.2 Analysis I: Varying *α*

Our first analysis with TINNiK used a range of *α* values for the *T* 3 test to detect quartet hybridization, but set *β* = 1 which, in effect, treats all quartets as resolved. Approximate ranges of *α* for which the full tree of blobs and individual blobs are detected are shown in Table 2.

With sample sizes *n ≥* 500 gene trees and ILS low or moderate, the tree of blobs is correctly inferred for a wide range of *α*. With 1000 gene trees sampled under low ILS condition, for example, the tree of blobs is correctly inferred for *α* ranging over eleven orders of magnitude. Even with high ILS, TINNiK returns the true tree of blobs from sample size 10,000.

A typical pattern of increasing resolution in the TINNiK tree of blobs as test level *α* is varied is shown in Figure 3, for *n* = 1000 and *k* = 1. Smaller *α* sets a stricter criterion for a quartet to be judged non-tree-like, so the count of quartets initially flagged as B-quartets in the algorithm is decreased, and the number of B-quartets inferred using the inference rule of Theorem 2 may shrink as well. Proceeding from large *α* to small,

a. TINNiK first detects the tree-like *A*-group,
b. the *C*-blob is detected, and a cut edge separating the {*C*-blob, *A*-group} from the *B*- and *D*-groups then appears,
c. the *D*-blob and then the *B*-blob are detected, so that the full tree of blobs is inferred for a range of *α* ∈ [10^*−*7^, 0.001],
d. the *B*- and *D*-blobs become increasingly over-resolved, although the *A*-group and *C*-blob are correctly inferred even for very small values of *α*.

Several additional patterns from Table 2 and Figure 3 hold in a wide range of our experiments. First, detecting features of the tree of blobs by TINNiK is harder for some parts than others. For instance, decreasing *α*, the *C*-blob and the *A*-tree group are the first parts to be correctly detected by TINNiK, and remain correctly resolved for a large range of test levels. This suggests that the metric structure and topological complexity of a network may result in varying difficulty in correctly inferring specific parts of the tree of blobs. A single analysis may be insufficient to explore all hybridization in a large network.

Second, when ILS is present in anything other than low amounts, a gene tree sample of size 300 drawn from *𝒩*^+^ appears too small to correctly infer the tree of blobs by TINNiK. Empiricists should be aware that the number of genes needed for accurate hybridization detection may be large. Whether these observations apply more generally to other data types and inference frameworks is unknown, as other tractable inference methods for non-level-1 networks are not yet available.

#### 5.2.3 Analysis II: Varying *β*

Short branches on a network result in higher levels of ILS, which can cause *CF* s to be closer to (1*/*3, 1*/*3, 1*/*3). To study these effects on TINNiK’s inference, a second analysis focused on the network *𝒩*^+^ of Figure 1 (L) with *k* = 0.5. We fixed *α* = 10^*−*4^ and varied the level *β* for the star tree test. Under this test, as *β* is decreased more quartets are taken to be star trees initially (and flagged as B-quartets) leading to more polytomies and less resolution in the TINNiK tree of blobs.

Figure 4 shows results for a sample size of *n* = 1000 gene trees. Proceeding from left to right, we see that for many values, *β >* 10^*−*6^, the *A*-group and *C*-blob are correctly detected. As *β* is decreased, the *A*-, *C*-, and *D*-groups are correctly inferred, then the correct tree of blobs is found for *β* ∈ [10^*−*34^, 10^*−*9^]. Decreasing *β* further results in the *D*-blob collapsing incorrectly (for instance, in the rightmost tree of Figure 4, {*D*2, *D*3} no longer form a cherry), and ultimately a star tree is produced.

Figure 5 shows typical simplex plots displaying the results of hypothesis tests (L) and application of the inference rule (R), for TINNiK test levels producing the true tree of blobs. The TINNiK algorithm first finds B-quartets corresponding to the red, green, and gold symbols displayed on the left. The increase in the number of B-quartets from the inference rule is visible in the gold symbols on the right.

That TINNiK can correctly infer the true tree of blobs when some *B*-quartets are found from the star tree test should be contrasted with results shown in Table 2 for this simulated data, where without *B*-quartets from the star tree test the tree of blobs was never correctly inferred. Using the star tree test to judge more quartets as unresolved (decreasing *β*) can thus help in obtaining the correct tree of blobs. High amounts of ILS from short branches can have the same qualitative impact on *CF* s as some blob structures, tending to equalize the entries, so that they are close to (1*/*3, 1*/*3, 1*/*3). The star tree test, by flagging such quartets as B-quartets regardless of the cause, helps prevent spurious resolution not strongly supported by the data.

#### 5.2.4 Analysis III: Varying blob complexity

To investigate the effect that blob complexity might have on TINNiK’s inference we considered a level-1 network 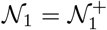 with a single 7-cycle, and then modified it by adding two additional hybridizations resulting in a level-3 network 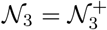. Figure 6 (L) shows *𝒩*_3_, with *𝒩*_1_ composed of only the black and magenta edges. A simulated sample of *n* = 1000 gene trees was analyzed with TINNiK.

For all ^10^ 4-taxon sets the expected quartet concordance factors were computed with `QuartetNetworkGoodnessFit` [39] and plotted in Fig. 6 (R, top) (*𝒩*_1_ magenta, *𝒩*_3_ blue). Expected *CF* s not on the model lines in Fig. 2 correspond to B-quartets, while some of those on the model lines may be inferred as B-quartets using the inference rule. The effect of increasing topological complexity in this 7-blob is to “pull” many *CF* s closer to the centroid.

The pull of *CF* s toward the centroid with increasing topological complexity means that the signal for hybridization increasingly resembles that for lack of quartet resolution. Intuition for this is that each particular choice of lineage paths through a blob determines a *CF* in the simplex, with a convex sum of these giving the expected *CF*. But a convex sum of a collection of *CF* s will be their weighted center of gravity, and hence tend toward their “middle.”

Since *CF* s computed from inferred gene trees in simulation studies have also been observed to be pulled toward the centroid from their expectation [27], the blurring of hybridization signal and lack of resolution may be very difficult to untangle. This suggests there may be practical limits on how complicated blob structure can be for reliable inference from *CF* s. Similarly, for fixed *α* = 10^*−*25^, TINNiK infers the true tree of blobs for a much wider range of *β* levels for the level-3 network *𝒩*_3_.

Table 3 shows a range of values for which the tree of blobs is correctly inferred when only one of the two test levels is varied. When *β* = 1, TINNiK infers the true tree of blobs for a much wider range of test levels *α* for *𝒩*_1_ than *𝒩*_3_. This is not surprising, since more of *𝒩*_1_’s *CF* s are placed distant from the model lines than those for *𝒩*_3_.

**Table 3.** Range of *α, β* values, for blob detection in 7-blob networks for a sample of 1000 gene trees.

| | $T3$ test:<br>$\alpha$ interval, $\beta = 1$ | star tree test:<br>$\alpha = 10^{-25}$, $\beta$ interval |
| --- | --- | --- |
| $\mathcal{N}_1$ | $[2 \times 10^{-10}, 0.03]$ | $[1 \times 10^{-83}, 1 \times 10^{-13}]$ |
| $\mathcal{N}_3$ | $[0.002, 0.01]$ | $[1 \times 10^{-108}, 1 \times 10^{-7}]$ |

#### 5.2.5 Analysis IV: Comparison to network inference

Some recent network inference methods seek to infer a level-1 network, yet offer no means of testing that assumption. One way that TINNiK might be helpful for this is by comparing its tree of blobs to an inferred level-1 structure. To test this possibility, we considered the level-2 network *𝒩*^+^ of Figure 7 (L), whose tree of blobs is a star tree. We analyzed simulated data of *n* = 10, 000 gene trees from this network using SNaQ [37], which assumes the network is level-1. We thus knowingly violated SNaQ’s assumptions, and did not expect its output to necessarily resemble the true network. SNaQ’s optimal level-1 networks with 1 and 2 hybridizations are shown in Figure 7 (C,R). Note that the network 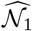 returned by SNaQ when *h*_*max*_ = 1 can not be obtained from *𝒩*^+^ by removing a single hybrid edge, nor is its tree of blobs 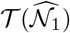 a star tree. Much of the inferred metric information also has little relationship to the true network’s branch lengths. When *h*_*max*_ = 2, the inferred SNaQ network 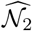 has two cycles joined with a branch of length zero. While the tree of blobs for 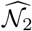 would be a star tree if the zero branch length were collapsed, the inferred blob structure is misleading. For instance, the close hybrid relationship between *D* and *E* is inferred as a more distant non-hybrid one.

The tree of blobs inferred by TINNiK is a (correct) star tree for any *α >* 10^*−*199^ and *β >* 10^*−*109^. For no values of *α* does TINNiK obtain a tree of blobs reflecting any of the individual cycles that SNaQ infers. Since both SNaQ and TINNiK base their inference on the same quartet *CF* s, the conflict is even more striking.

### 5.3 Empirical data

We apply TINNiK to infer trees of blobs from several empirical datasets: Hawaiian flowering plants [40] and primates [38]. These have been analyzed for hybridization previously, with conflicting results depending on the method used.

#### 5.3.1 Hawaiian *Cyrtandra*

A recent study by Kleinskopf et. al. [40] investigated hybridization and introgression in the Hawaiian *Cyrtandra*. Although samples were collected across the islands, network analyses by PhyloNet and SNaQ were restricted to single island subsamples. The dataset consists of 569 gene trees, a few with missing taxa. Most of the gene trees are poorly resolved, with a majority of gene quartet trees unresolved for almost all sets of 4 taxa. For the Kauai island group of 7 taxa, for example, the star tree topology is a majority for all but one gene quartet (34 of 35).

Networks inferred by PhyloNet and SNaQ (with *h*_*max*_ = 1) for the Kauai group agreed [40, Fig. 4]. Since the lack of gene tree resolution indicated information content might be low, TINNiK analyses were performed both with unresolved gene quartets omitted, and with unresolved quartets apportioned uniformly among the three resolved topologies. TINNiK’s analyses support the PhyloNet/SNaQ underlying tree of blobs (Figure 8 (L)) over a wide range of test levels with both methods for handling unresolved gene quartets. Specifically, when unresolved quartets are included in the analysis, the supporting TINNiK tree of blobs shown in Figure 8 (L) is obtained for any *α* ∈ [0.03, 0.14] with *β* = 1, and for *α* = 0.05 with *β*∈ [0.3, 1].

**Fig. 8.**
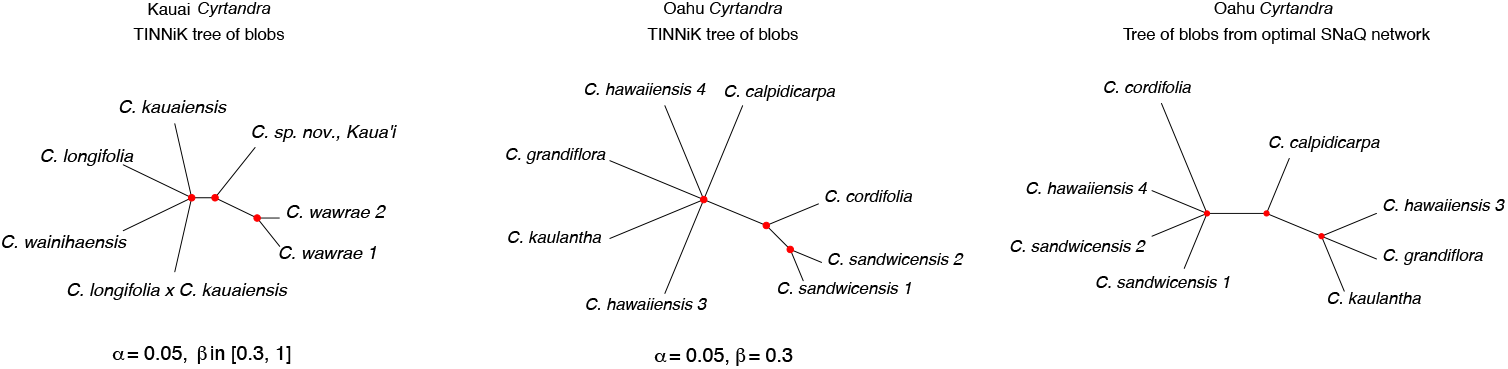
(L) For the Kauai dataset, TINNiK’s tree of blobs supports the PhyloNet and SNaQ analyses for a wide range of test levels *α, β*, regardless of handling of unresolved quartet topologies. (C) The TINNiK tree of blobs for the Oahu dataset when *α* = 0.05 and *β* = 0.3 conflicts with both SNaQ and PhyloNet analyses. It supports some blob structure, but not that inferred by SNaQ. (R) The tree obtained by contracting cycles in the SNaQ network inferred from the Oahu dataset. PhyloNet infers a resolved tree for these data.

In contrast, there is considerable discrepancy between the PhyloNet and SNaQ analyses for the 8-species Oahu group [40]. PhyloNet infers a tree, while SNaQ infers a level-1 network with 2 cycles. We found that TINNiK inferred exactly three topologies as *α* and *β* were varied: 1) a binary tree agreeing with that inferred by PhyloNet, 2) a tree of blobs 𝒯 pictured in Figure 8 (C) with exactly 2 cut edges, and 3) a star tree. When *α* is small and *β* large, so that there are no initial B-quartets, the inferred TINNiK tree agrees with that of PhyloNet (and `MSCquartets`’ QDC tree [6]). This supports PhyloNet’s analysis in that signal for hybridization in the Oahu data may be weak. Moreover, the inferred TINNiK tree 𝒯 of Figure 8 (C) does not agree with the tree of Figure 8 (R) obtained by contracting cycles in SNaQ’s optimal network. Possible reasons for this conflict might again be signal too weak for these analyses, or that the underlying network is not level-1. Regardless of the cause, TINNiK illuminates that further investigation is needed to understand relationships in this group.

#### 5.3.2 Primate data

A recent study of primates by Vanderpool et al. [38] used full genome data to investigate phylogenetic relationships between 26 primates. Multiple analyses were performed, but we focus on two investigations, into resolution of a clade of New World Monkeys (NWMs), and of possible introgression within a subset of 7 taxa, the Papionini group. These data were also studied in [18] using PhyNEST. Input for our analyses were the 1730 gene trees estimated in [38].

The placement on the primate tree of some NWMs is uncertain, with one analysis supporting that *A. nancymaae* and *C. jacchus* form a clade sister to the {*S. boliviensis, C. Capucinis imitator*}clade, and a second that *A. nancymaae* is sister to the {*S. boliviensis, C. Capucinis imitator*}clade with *C. jacchus* an outgroup [38]. Using `MSCquartets` to compute empirical *CF* s and to perform hypothesis tests, we found that quartet *CF* s that clustered near the centroid are exactly those that might resolve this issue. In Figure 9 (L) for any *β <* 0.1 (shown with *α* = 10^*−*7^), the golden squares clustered around the centroid where the star tree hypothesis is not rejected for any alternative resolved topology, are those involving {*C. jacchus, A. nancymaae*}, exactly one of {*S. boliviensis, C. capucinus imitator*} and a fourth taxon. As seen in Figure 9 (R), the TINNiK tree of blobs has a degree 4 node for any *β <* 0.1, which does not support further resolution.

**Fig. 9.**
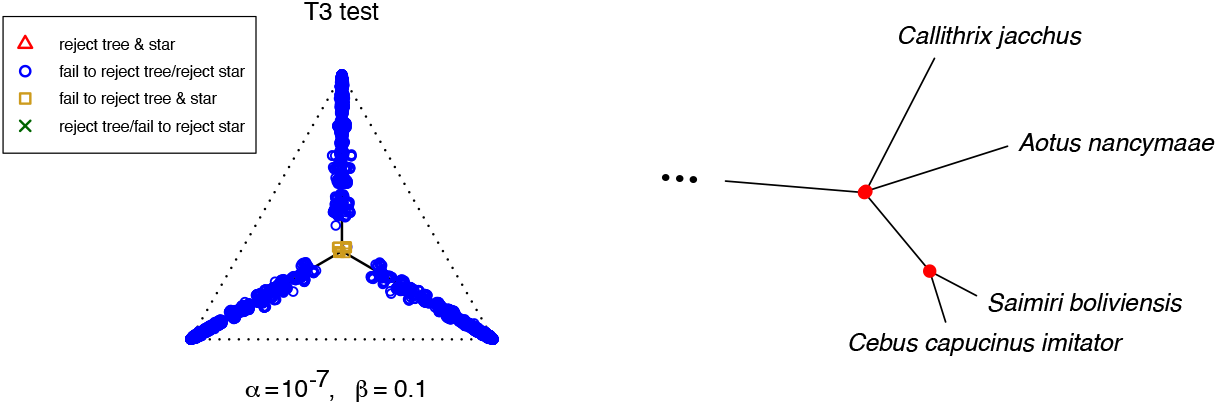
(L) Simplex plots illustrate that hypothesis test results support the star tree topology for quartets with *A. nancymaae, C. jacchus* and one of the other two NWMs, and (R) close up of tree of blobs for the NWMs for any *β ≤* 0.1.

A subset of four Asian Papionini (*Cercocebus atys, Mandrillus leucophaeus, Papio anubis, Theropithecus gelada*) and three African Papionini (*Macaca ascicularis, Macaca mulatta, Macaca nemestrina*) were also analyzed by Vanderpool et. al., with multiple introgression events found between and among these groups [38, Fig. 4] using the Δ method of [41]. Specifically, seven introgression events were inferred, with four crossing continental boundaries.

The sequence data for these taxa were reanalyzed by Kong et. al. using PhyNEST to infer level-1 networks with *h*_*max*_ = 1, 2 hybridizations [18]. *Theropithecus gelada* was found to be a hybrid of *Papio anubis* and *Mandrillus leucophaeus* when *h*_*max*_ = 1, with an additional hybridization among the Macaques when *h*_*max*_ = 2. These level-1 hybridization cycles do not cross continental boundaries.

A TINNiK analysis of these seven taxa was performed with *β* = 1, since the simplex plot of Figure 10 (L) shows no *CF* s close to that of the star tree. In Figure 10 (C) the TINNiK tree of blobs for test levels *α* ∈ [0.008, 0.032] agrees with the PhyNEST analysis when *h*_*max*_ = 1. For larger *α*∈ [0.033, 0.093] the tree of blobs is shown (R), with multifurcations for each continent, consistent with the PhyNEST analysis when *h*_*max*_ = 2. For test levels *α >* 0.093, TINNiK returns the star tree, consistent with the analysis using Δ. However, such a large value of alpha indicates weak support for additional hybridization spanning continents.

**Fig. 10.**
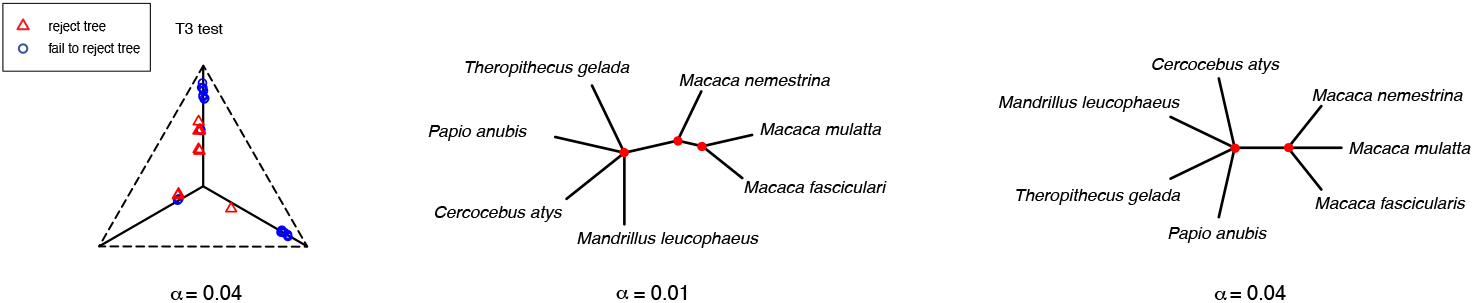
(L) Results of *T* 3 hypothesis tests for *α* = 0.04, *β* = 1; (C) For *α* ∈ [0.008, 0.032] TINNiK’s tree of blobs supports hybridizaton among the African Papionini; and (R) For *α* ∈ [0.033, 0.093] TINNiK supports hybridizaton within the African Papionini and the Asian macaques. Only for larger values of *α* does TINNiK return a star tree of blobs.

## 6 Conclusions

The implementation of the TINNiK algorithm in `MSCquartets` provides the first software tool for statistically-justified inference of the tree-like parts of a species network. With input of gene trees inferred from multilocus sequence data, it returns an inferred tree of blobs of the network under the NMSC, without restrictive assumptions on the reticulation structure within the blobs.

In some cases, the tree of blobs, perhaps with partial information on individual blob structure, may represent the most we can tell about a species network from biological data. While the theoretical limits to inference of complex blob structure are still unknown, recent work [19] has shown that different blob structures are indistinguishable from certain types of commonly-used gene tree summary data. Even in cases where theoretical identifiability holds, practical identifiability may not, as the signal distinguishing the precise structure may be obscured by even small levels of noise. Learning a blob is present, or only part of its structure, may be the strongest practical inference that can be performed for some data.

When more can be inferred, the tree of blobs for a large group of species can provide a good starting point for a more targeted investigation into the unknown relationships represented by its multifurcations. Its inference might be either a first step in an exploratory data analysis, or form the basis for a divide-and-conquer approach, although the more demanding statistical inference of internal blob structure requires further theoretical and practical development.

Several recently-proposed network inference methods assume a level-1 structure (SNaQ, PhyNEST, NANUQ), but their performance under level misspecification has not been studied in published work. (Note, however, that NANUQ’s splits graph can suggest such model violation.) Since TINNiK does not assume a particular level or other special blob structure, it can provide an important alternative perspective. Inference of a full network that is incompatible with TINNiK’s tree of blobs can suggest possible model violations and a need for further analysis. If gene trees have already been inferred, TINNiK’s speed means its use in this way requires little additional computational effort.

An inferred tree of blobs may also be useful for the heuristic searches performed by methods attempting to find complete networks. When a starting network is needed, a TINNiK tree of blobs is a natural candidate so the search may spend less time finding the tree-like parts of the network. Even if TINNiK produces an over-resolved tree, in our experiments this is often a tree displayed on the network, so that as new hybrid edges are introduced the search may still soon focus on good candidate networks. Finally, for those methods requiring an *a priori* upper bound on the number of reticulations, TINNiK can again be helpful by suggesting the number of blobs, and thus a minimum number of reticulations needed.

Although our justification of the TINNiK algorithm in this work has emphasized the NMSC model, its essential ideas could be applied to other models of gene tree formation. For instance, recent work [19] considered two other models, one in which gene trees must be displayed on the species network so coalescence is immediate, and a common-inheritance coalescent model in which the standard coalescent applies but only inside displayed trees. In both these cases it is possible to identify B-quartets for 4-taxon networks from certain data types, and thus follow the outline of our algorithm.

The introduction of TINNiK for inferring the tree of blobs of a species network from biological data should encourage the development of other algorithms for this problem. Network inference remains difficult for both theoretical and practical reasons, and phylogenomics will benefit from an expanding array of approaches.

## Acknowledgements

We thank Kristina Wicke for many helpful conversations, and for testing the software implementation extensively.

## Declarations

### Code availability

TINNiK is implemented as part of the `MSCquartets` 2.0 R package, freely available on CRAN.

### Data availability

Empirical data, all generated by other researchers in previous publications, are publicly available.

## Appendix A Newick for model networks

The Newick string for the model network shown in Figure 1 (L) used in simulations in Sections 5.2.2 and 5.2.3 is:

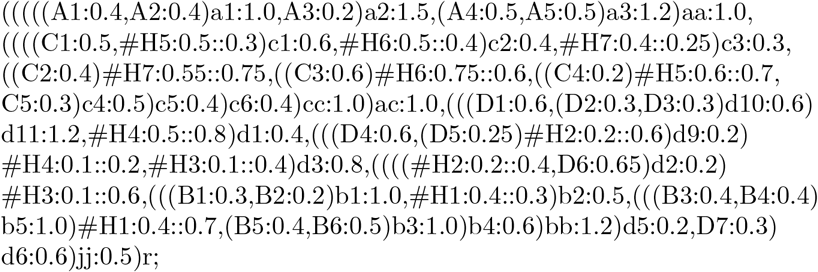

The Newick string for the model network shown in Figure 6 (L) used in of Section 5.2.4 is:

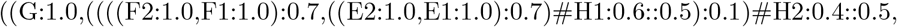

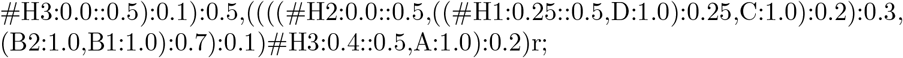

The Newick string for the model network shown in Figure 7 (L) used in of Section 5.2.5 is:

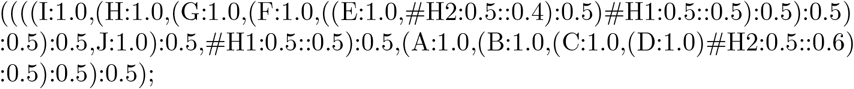

## Appendix B B- and T-quartets on a sunlet network

**Fig. B1.**
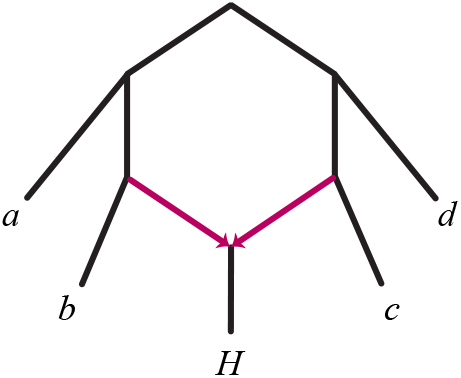
The quartet *Q* = {*a, b, c, d*,} is a B-quartet on *𝒩* ^+^, but a *T*-quartet on the induced network 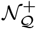

## Appendix C Cut hypothesis test and simulations

We use the notation of Section 3.1.

### C.1 Cut model topology estimation and LR statistic

For the cut model, the maximum likelihood parameter estimate for data

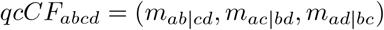

with *m* = *m*_*ab*|*cd*_ +*m*_*ac*|*bd*_ +*m*_*ad*|*bc*_ is found by computing the maximum of the trinomial likelihood constrained to each of the 3 line segments of Figure 2 (L) and then choosing the largest. (Ties broken at random.) For the vertical line the maximizer is

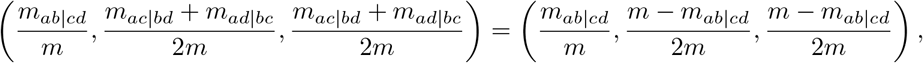

the projection of 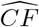 (the normalized *qcCF*) orthogonally to the line. A comparison of the likelihood at the maximizers on the three lines leads to the three regions shown in Figure C2 for which normalized *qcCF* s lead to cut model MLEs on the model lines in each region. For use in TINNiK, when the cut model is not rejected, we need only the topology of the MLE, which is determined solely by the color of the region in which 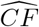 lies.

**Fig. C2.**
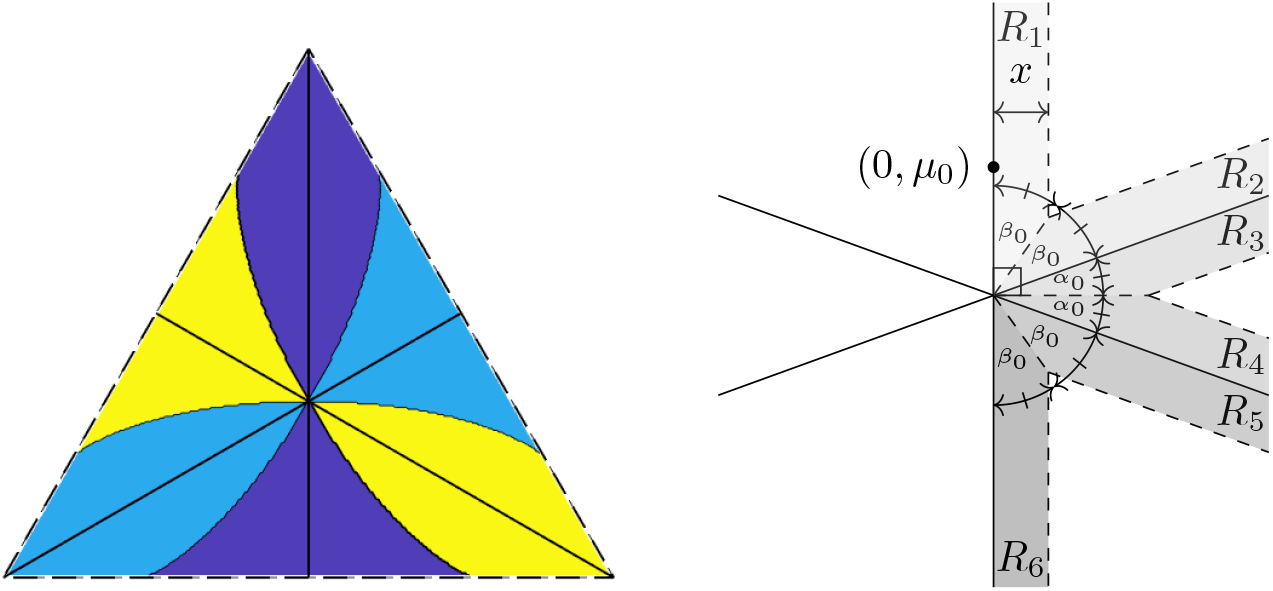
(L) Regions for which data with an empirical 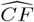 gives an MLE in the cut model on each of the three cut model lines. The MLE is obtained by moving orthogonally to the model line in the same colored region as 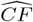. (R) The image of the cut model in Δ_2_ under a linear transformation to the plane, with 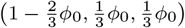 mapped to (0, *µ*_0_), and the 3 model lines mapped to the three lines shown. The region of integration for *G* (*x*) in the proof of Proposition 6 is all shaded regions *R*_*i*_ and their reflections about the vertical line.

The likelihood ratio statistic for the hypothesis test described in Section 3.1 requires the maximum log-likelihoods under the cut (null) and unconstrained (alternative) trinomial models. For the vertical line of the cut model, the maximum log-likelihood is

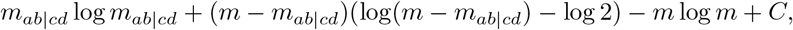

while for the unconstrained model, with MLE 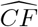, the maximum log-likelihood is

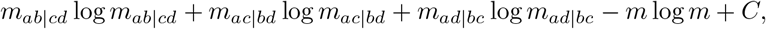

with *C* a constant.

### C.2 Cut test distributions

To derive an asymptotic distribution for the likelihood ratio (LR) statistic for the cut model hypothesis test of Section 3.1, we follow similar derivations for *T* 1 and *T* 3 tests using Theorem 3.1 of [28]. That work also provides more discussion of why model singularities, such as the cut model’s (1*/*3, 1*/*3, 1*/*3), make the use of a standard distribution inappropriate.

Assume the generating parameter in the cut model is *θ*_0_ = (1 *−* 2*φ*_0_*/*3, *φ*_0_*/*3, *φ*_0_*/*3). Applying a linear transformation dependent on the sample size *n* and the Fisher information matrix *ℐ* as in [28], the simplex is mapped to ℝ^2^, and the cut model to lines crossing at the origin, with the vertical line segment mapped to the *y*-axis. Then *θ*_0_ *↦* (0, *µ*_0_), with 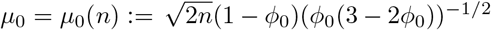, and the information matrix becomes the identity. This transformation is not conformal (unless *φ*_0_ = 1) and the two transformed model lines not containing the generating parameter form angles *α*_0_ = arctan (3(3 *−* 2*φ*_0_))^*−*1*/*2^ with the horizontal. See Figure C2 (R).

#### Proposition 5.

*The likelihood ratio statistic for testing H*_0_ *versus H*_1_ *for a true parameter point θ*_0_ = (1*−* 2*φ*_0_*/*3, *φ*_0_*/*3, *φ*_0_*/*3), *φ*_0_∈ (0, 3*/*2), *of the cut model, with sample size n is asymptotically distributed as the random variable*

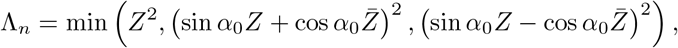

*where* 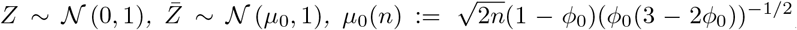, *and α*_0_ = arctan. (3(3 *−* 2*φ*_0_))^*−*1*/*2^

*Here”asymptotically distributed” means that the likelihood ratio statistic and this random variable converge in distribution to the same limit as n → ∞*.

*Proof*. Letting *γ*_0_ = tan *α*_0_, in the transformed space the image of Θ_0_ is contained in the union of the lines *x* = 0, *y* = *γ*_0_*x* and *y* = *− γ*_0_*x*.

By Theorem 3.1 of [28], the approximate distribution of the likelihood ratio statistic is the distribution of the minimum squared Euclidean distance between a normal sample, *𝒩* ((0, *µ*_0_), *I*), and the three lines in the transformed space. Assuming that *θ*_0_ is not too close to the boundary of the simplex, in a sense dependent on the sample size, little of the mass of *𝒩* ((0, *µ*_0_), *I*) is outside the image of the simplex. Thus, for the remainder of the argument, we replace these line segments with lines intersecting at the singularity (0, 0).

Denote the marginal probability distributions of the bivariate normal sample by *Z ∼ 𝒩* (0, 1) and 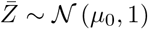. We next determine the squared distance of a sample point 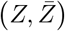 to each of the three lines.

Considering the first line, *x* = 0, the squared Euclidean distance is *Z*^2^. For the line *y* = *γ*_0_*x*, the closest point (*X, γ*_0_*X*) to 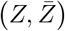 has 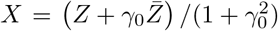, so the squared distance is

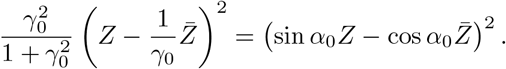

Similarly, for the line *y* = *−γ*_0_*x* the squared distance is 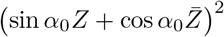. The claim follows by taking the minimum of these squared distances.

For testing purposes, we characterize this distribution further.

#### Proposition 6.

*The probability density function for the random variable* 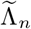 *of Proposition 5 is, for λ >* 0,

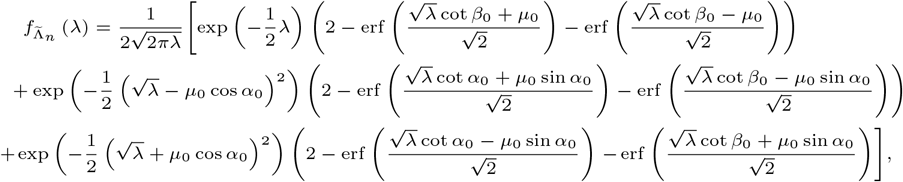

*where α*_0_ = arctan ((3(3 *−* 2*φ*_0_))^*−*1*/*2^) *and* 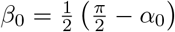.

*Proof*. To determine the probability density function for the distribution of Proposition 5, let *G* (*x*) denote the cumulative distribution function of the (non-squared) Euclidean distance. This is found by integrating the distribution *𝒩* ((0, *µ*_0_), *I*) over the tube of points within distance *x* from the image of Θ_0_, with simplifications using the symmetry of the region and normal distribution as shown in Figure C2 (R). Although the generating parameter (0, *µ*_0_) is shown above the origin, for *φ*_0_ ∈ 1, ^3^ it may be below. In fact, *α*_0_ is an increasing function of *φ*_0_, with *α*_0_ (0) *≈* 0.322 and 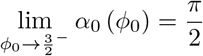.

Then 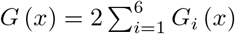, where *G*_*i*_ is the integral over the shaded strip *R*_*i*_, and the density of the Euclidean distance is 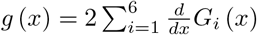.

Considering 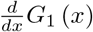 first:

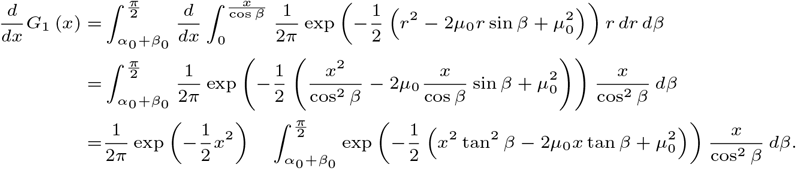

Substituting *y* = *x* tan *β* gives

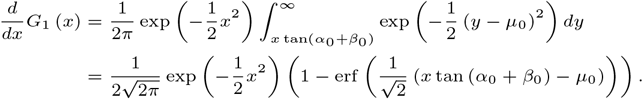

Next we consider 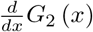 and 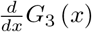. Rotating the figure 2*β*_0_ counter-clockwise to simplify the computation, the generating parameter is mapped as (0, *µ*_0_) *↦* (*−µ*_0_ cos *α*_0_, *µ*_0_ sin *α*_0_). Then

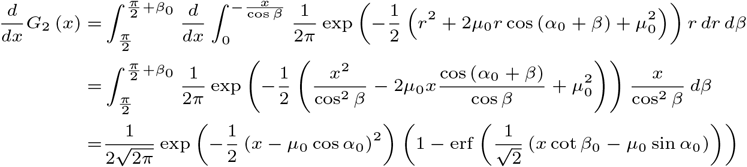

and

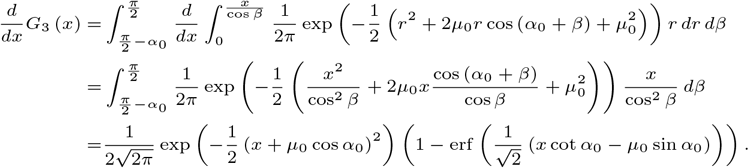

Next we consider 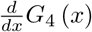 and 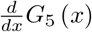. Rotating the figure 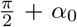 counter-clockwise, the generating parameter becomes (*−µ*_0_ cos *α*_0_, *−µ*_0_ sin *α*_0_). Then

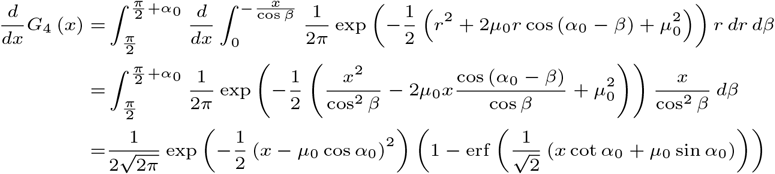

and

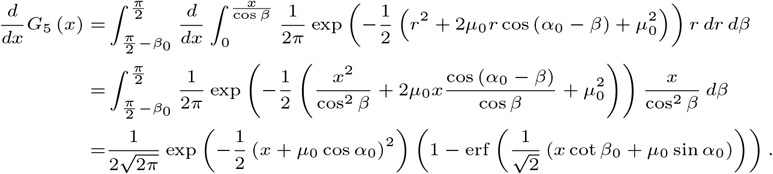

Finally, we consider 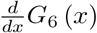, which is identical to 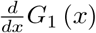, but after mapping the generating parameter as (0, *µ*_0_) *↦* (0, *−µ*_0_). Then

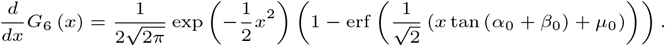

The claim follows after noting that tan (*α*_0_ + *β*_0_) = cot *β*_0_ and performing a change of variable to the squared Euclidean distance.

Using the density of Proposition 6 for judging likelihood ratio statistics is still complicated by its dependence on the unknown true parameter *φ*_0_. While several approaches to deal with this are discussed in [28], the simplest is to replace *φ*_0_ by its MLE under the cut model. Although theory does not guarantee the good performance of this approach, in the next section we investigate performance through simulation.

If an expected *qcCF* has some small counts, say less than 5, and hence its normalization lies near the boundary of the simplex, an alternative testing procedure is necessary. In the case of only 2 small counts, the geometry of the parameter space far from a vertex can be ignored, giving essentially the same situation for the *T* 1 and *T* 3 tests already implemented in `MSCquartets`. Either parametric bootstrapping from 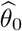, or a much faster precomputed approximation, can be used.

If only one expected count is small, the normalization lies near an edge of the simplex. Under the cut model, the other two counts should be approximately equal and approximately binomially distributed. Then a standard binomial test can be applied.

#### C.3 Cut test simulation

Figure C3 shows results of simulations comparing *p*-values for the likelihood ratio statistic for simulated *qcCFs* from the cut model, using the distribution of Proposition 6 with the MLE for *φ*_0_, and a standard 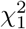 distribution. While neither distribution produces the desired cumulative distribution of *p*-values, that of Proposition 6 comes closer when *µ*_0_ has smaller magnitude. The value *µ*_0_ = 0 corresponds to the model singularity (1*/*3, 1*/*3, 1*/*3), where both tests will perform very conservatively for small significance levels, seldom rejecting the null model. As *µ*_0_ is varied away from 0, performance improves.

While *µ*_0_ depends on both the true model parameter and the sample size *n*, it has a simple interpretation: if *σ*_*y*_ is the standard deviation of the *y*-coordinate of the random observations, then |*µ*_0_|*σ*_*y*_ is the distance between the generating parameter *θ*_0_ and the model singularity.

**Fig. C3.**
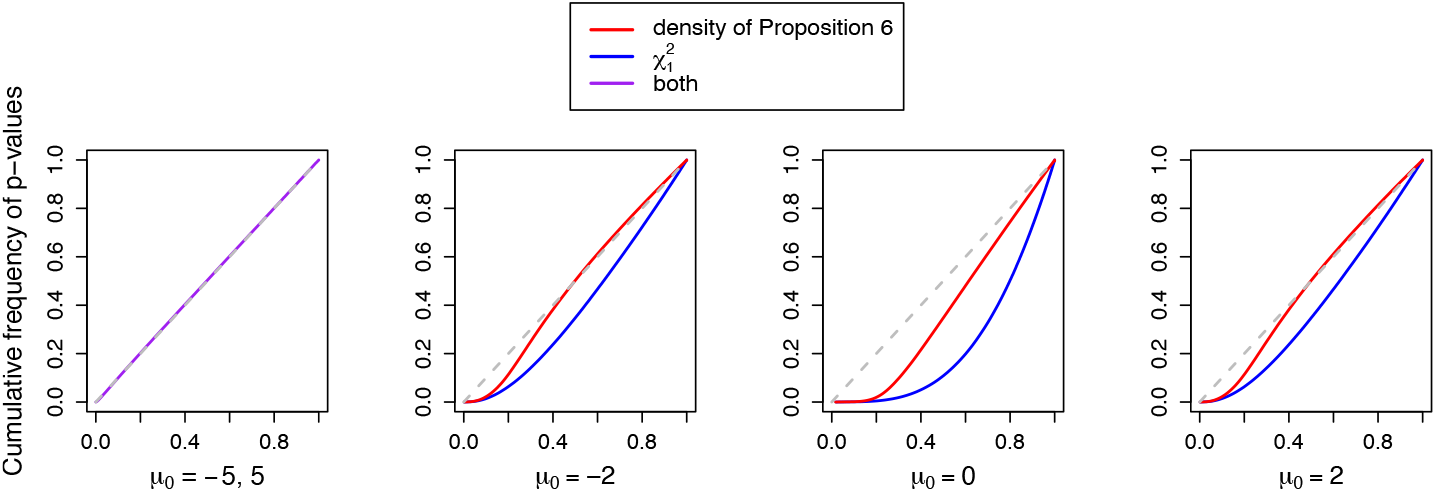
Cumulative distributions of *p*-values computed from simulations, for the distribution of the likelihood ratio statistic given in Proposition 6 using maximum likelihood estimates of *φ*_0_ (red), and for the 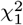 distribution (blue) for sample size *n* = 10^6^. The cdf plots are indistinguishable for |µ_0_| large. The diagonal line represents ideal behavior. At and near the model singularity, *µ*_0_ = 0, the distribution of Theorem 6 performs better than a 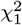.

## Footnotes

1 ”Tinnik” is the Inupiaq word for bearberry or kinnickinnick, a ground plant found throughout the circumpolar north.

